# The versican-hyaluronan complex provides an essential extracellular matrix niche for Flk1+ hematoendothelial progenitors

**DOI:** 10.1101/753418

**Authors:** Sumeda Nandadasa, Anna O’Donnell, Ayako Murao, Yu Yamaguchi, Ronald J. Midura, Lorin Olson, Suneel S. Apte

## Abstract

Little is known about extracellular matrix (ECM) contributions to formation of the earliest cell lineages in the embryo. Here, we show that the proteoglycan versican and glycosaminoglycan hyaluronan are associated with emerging Flk1^+^ hematoendothelial progenitors at gastrulation. The mouse versican mutant *Vcan*^hdf^ lacks yolk sac vasculature, with attenuated yolk sac hematopoiesis. CRISPR/Cas9-mediated *Vcan* inactivation in mouse embryonic stem cells reduced vascular endothelial and hematopoietic differentiation in embryoid bodies, which generated fewer blood colonies, and had an impaired angiogenic response to VEGF_165_. HA was severely depleted in *Vcan*^hdf^ embryos, with corresponding increase in the HA-depolymerase TMEM2. Conversely, HA-deficient mouse embryos also had vasculogenic suppression but with increased versican proteolysis. VEGF_165_ and Indian hedgehog, crucial vasculogenic factors, utilized the versican-HA matrix, specifically versican chondroitin sulfate chains, for binding. Versican-HA ECM is an obligate requirement for vasculogenesis and primitive hematopoiesis, acts as an vasculogenic factor-enriching microniche for Flk1^+^ progenitors from their origin at gastrulation.

## Introduction

The first blood vessels in amniotes form *de novo* in the yolk sac prior to initiation of hemodynamic forces, indicating an in situ differentiation driven by the local environment, which includes extracellular matrix (ECM)[1]. This vasculogenic process is inextricably linked to subsequent primitive hematopoiesis, i.e., formation of the first erythroid and myeloid cells [2], implying a close lineage relationship between the first vascular endothelial and blood cells. The relevant Flk1^+^ progenitor cells arise from mesodermal precursors at gastrulation [3], migrate from the primitive streak, and aggregate in blood islands in the proximal extra-embryonic yolk sac on the seventh day of gestation (E7) in mouse embryos [4, 5]. Several key transcription factors and soluble effectors of vasculogenesis/angiogenesis and hematopoiesis are known [1]. Although ECM and ECM-derived proteolytic fragments are recognized as angiogenesis regulators [6] and sulfated glycosaminoglycans can promote endothelial differentiation of mesenchymal stem cells [7], specific ECM proteoglycans that influence vasculogenesis are unknown. Another early cell lineage, primordial germ cells, are accompanied by a protective “traveling niche” of steel factor/stem cell factor-producing cells during migration to the gonads [8]. Whether a cell-associated ECM supports other early lineages such as hematoendothelial progenitors is unknown.

ECM surrounds all mammalian cells, forming a distinct pericellular matrix in some cell types, and the interstitial matrix of tissues. It influences cell behavior by modulating cell adhesion, migration and tissue mechanics, sequesters growth factors and cytokines, and its proteolysis can generate bioactive fragments. Proteoglycans, which are ECM and cell-surface molecules with one or more glycosaminoglycan chains covalently attached to a core protein, participate in all these mechanisms. For example, chondroitin sulfate proteoglycans (CSPGs) such as versican, which are bulky and inherently anti-adhesive, regulate focal adhesion formation and cell migration [9–11]. Heparan sulfate proteoglycans (HSPGs) sequester growth factors, including angiogenic growth factors such as VEGF_165_ and FGF2 through their HS chains and act as co-receptors in angiogenic signaling [12]. CSPGs can also bind VEGF_165_ and may overlap functionally with HSPGs in VEGF-induced sprouting angiogenesis [13], but physiological in vivo contexts for this regulatory activity remain unidentified for CSPGs.

Versican is a widely distributed CSPG that aggregates with the glycosaminoglycan hyaluronan (HA), through its N-terminal G1 domain [14–17]. Its C-terminal G3 domain binds to ECM components fibronectin, fibrillins and tenascins [18] and was shown to interact with VEGF_165_ in biochemical assays [19]. HA is anchored to cell-surface receptors such as CD44 and RHAMM as well as the membrane-localized hyaluronan synthases [20], providing a means by which versican can localize to the pericellular matrix. Pericellular versican was previously shown to modulate smooth muscle cell differentiation via regulation of cell adhesion [9, 21, 22]. Four versican isoforms (V0, V1-V3) arise from alternative splicing of large exons, numbered 7 and 8, encoding CS-bearing domains GAG*α* and GAG*β*, respectively [16, 23]. A versican insertional mutant mouse allele (*Vcan*^hdf^), which lacks all isoforms, demonstrated an essential role for versican in early cardiac development [24, 25]. *Has2* null embryos, lacking the major HA synthase, die by 10.5 days of gestation with similar cardiac defects as *Vcan*^hdf/hdf^ [26], consistent with the versican-HA complex being a major constituent of cardiac jelly. Here, upon identification of abnormal vasculature in *Vcan*^hdf/hdf^ yolk sac, we investigated its role in vasculogenesis and early hematopoiesis, with investigation of the key findings in embryos with inactivation of hyaluronan synthase (*Has*) genes. An intimate association of versican and HA with Flk1^+^ hematoendothelial progenitor cells from their origin at gastrulation is demonstrated to have profound significance for vasculogenesis and primitive hematopoiesis.

## Results

### *Vcan*^hdf/hdf^ yolk sacs lack a vascular plexus

*Vcan*^hdf/hdf^ embryos (Figure 1-Supplement 1) did not survive past E10.5, and were recognizable by consistently smaller size and dilated pericardial sac at E9.5 (Fig. 1A). Although major developmental milestones, including axial rotation of the embryo and initiation of cardiac contraction were completed by E9.5, *Vcan*^hdf/hdf^ yolk sacs and embryos were less vascular (Fig. 1B,C). Whole mount CD31 immunostaining revealed the lack of a vascular network in E9.5 *Vcan*^hdf/hdf^ yolk sacs, instead of which scattered vascular CD31+ cells were present (Fig. 1C).

**Figure 1.**
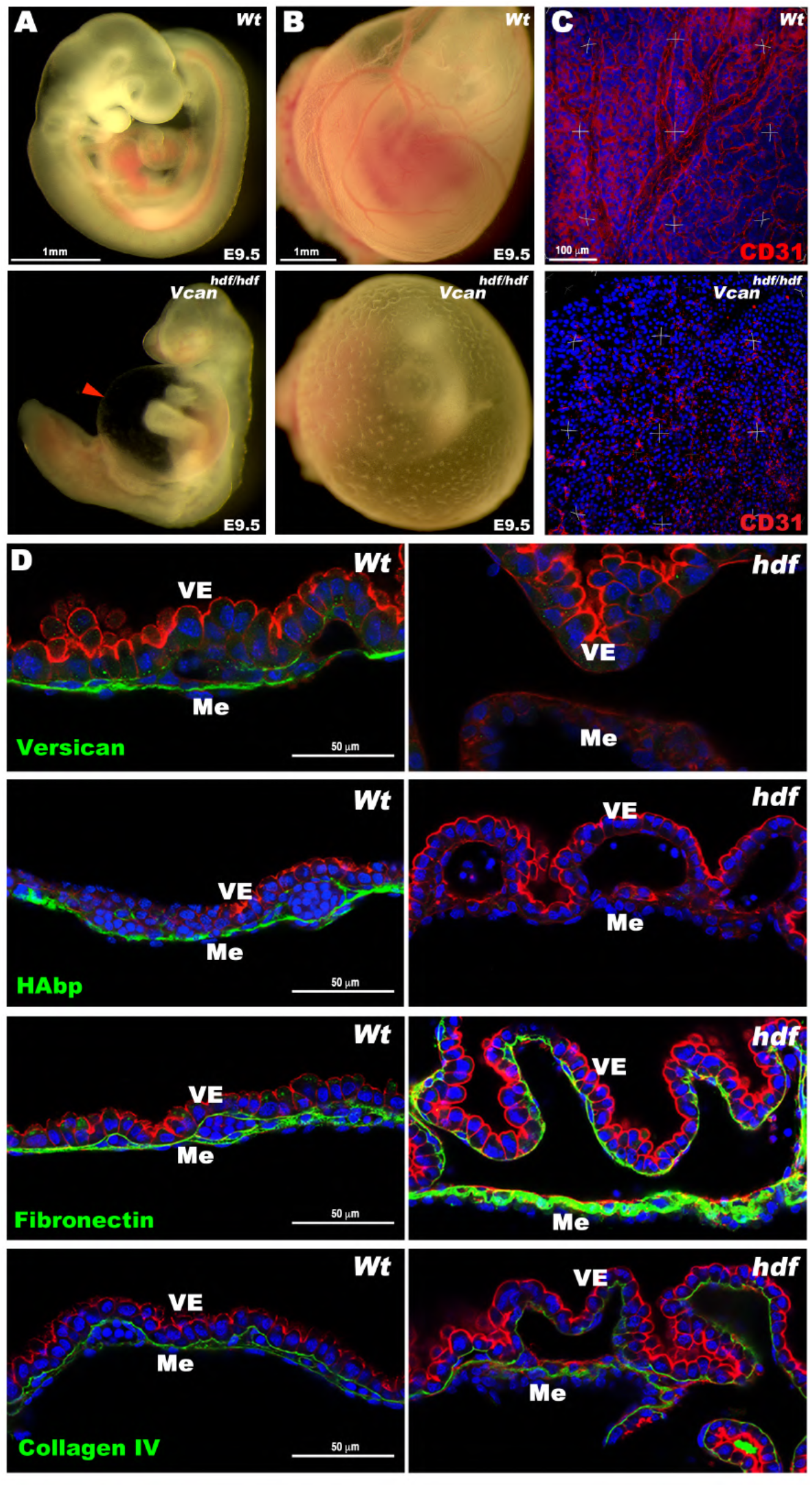
*Vcan^hdf/hdf^* yolk sacs are avascular and lack hyaluronan. **(A)** E9.5 wild type and *Vcan^hdf/hdf^* embryos. The red arrowhead shows the dilated pericardial sac in the latter. **(B)** E9.5 yolk sacs imaged in situ demonstrates the absence of vasculature in the E9.5 *Vcan^hdf/hdf^* yolk sac. **(C)** Three-dimensional (3D) maximum-intensity projections of whole-mount yolk sacs stained with anti-CD31 (red) showing absence of the vascular network in E9.5 *Vcan^hdf/hdf^* yolk sacs (n=3 yolk sacs of each genotype). **(D)** Versican GAG*β* staining (green) is present throughout the mesoderm (Me) of E9.5 wild type yolk sacs and absent in the *Vcan^hdf/hdf^* yolk sac. F-actin (red) staining highlights visceral endoderm (VE), detaches from mesoderm in *Vcan^hdf/hdf^* yolk sacs. HA and fibronectin (green) were similarly distributed as versican in wild type yolk sac, but HA staining was absent in *Vcan^hdf/hdf^* yolk sac and fibronectin staining was more intense. Collagen IV staining intensity was similar in wild type and *Vcan^hdf/hdf^* yolk sacs (n=3 yolk sacs of each genotype). Scale bar in A, B=1mm, C= 100μm, D= 50μm.

In E9.5 wild-type yolk sac versican localized to the mesoderm, but no staining was present in *Vcan*^hdf/hdf^ yolk sacs (Fig. 1D). HA-binding protein (HABP) staining overlapped with versican in wild-type yolk sac mesoderm but was absent in *Vcan*^hdf/hdf^ yolk sacs (Fig. 1D). Fibronectin immunostaining in contrast, showed increased staining in *Vcan*^hdf/hdf^ yolk sacs, whereas collagen IV staining was unaffected (Fig. 1D), suggesting that loss of HA did not reflect global ECM reduction in *Vcan*^hdf/hdf^ yolk sacs. Similar to *Has2*, *Itga5* and *Fn1*-null alleles, which have failed yolk sac vasculogenesis and hematopoiesis [26, 27], *Vcan*^hdf/hdf^ yolk sacs consistently showed mesoderm detachment from visceral endoderm with few evident blood islands (Fig 1D, Fig.1 Supplement 2 A-C). Intense F-actin staining was consistently observed in *Vcan*^hdf/hdf^ visceral endoderm (Fig. 1 D, Fig.1 Supplement 2A), and rounded, rather than spindle-shaped nuclei were observed in the yolk sac mesothelial layer (Fig.1 Supplement 2B). Transmission electron microscopy (TEM), undertaken with fixation conditions that preserved cell-matrix interactions [28] showed complete coverage of blood islands by vascular endothelium in wild-type yolk sac but discontinuous vascular endothelial cells with numerous membrane protrusions in *Vcan*^hdf/hdf^ yolk sacs (Fig.1 Supplement 2C). Thus, versican is essential for proper yolk sac morphogenesis.

E8.5 *Vcan*^hdf/hdf^ embryos had a normal shape, but appeared pale and their yolk sacs lacked a visible vascular network (Fig. 2A). Whole-mount CD31 immunostaining of E8.5 embryos identified severe, widespread attenuation of vasculature (Fig. 2B). *En face* imaging of whole-mount wild-type yolk sac stained with anti-CD31 demonstrated a well-formed vascular plexus lacking in *Vcan*^hdf/hdf^ yolk sacs (Fig. 2C). Immunostaining of versican and CD31, together with HA-staining in E8.5 wild-type yolk sacs revealed versican and HA co-localization in patches corresponding to individual cells on the mesothelial aspect of blood islands (Fig. 2D). mRNA *in situ* hybridization with *Vcan* exon 7 (GAG*α*), or exon 8 (GAG*β*)-specific probes revealed that only the exon 8 probe hybridized to E8.5 yolk sac mesoderm (Fig 2E), suggesting exclusively versican V1 isoform expression at E8.5. Both *Vcan* probes hybridized strongly to E8.5 wild-type hearts (Fig. 2F) but not *Vcan*^hdf/hdf^ hearts (e.g., exon 8 probe, Fig. 2G), indicative of specificity.

**Figure 2.**
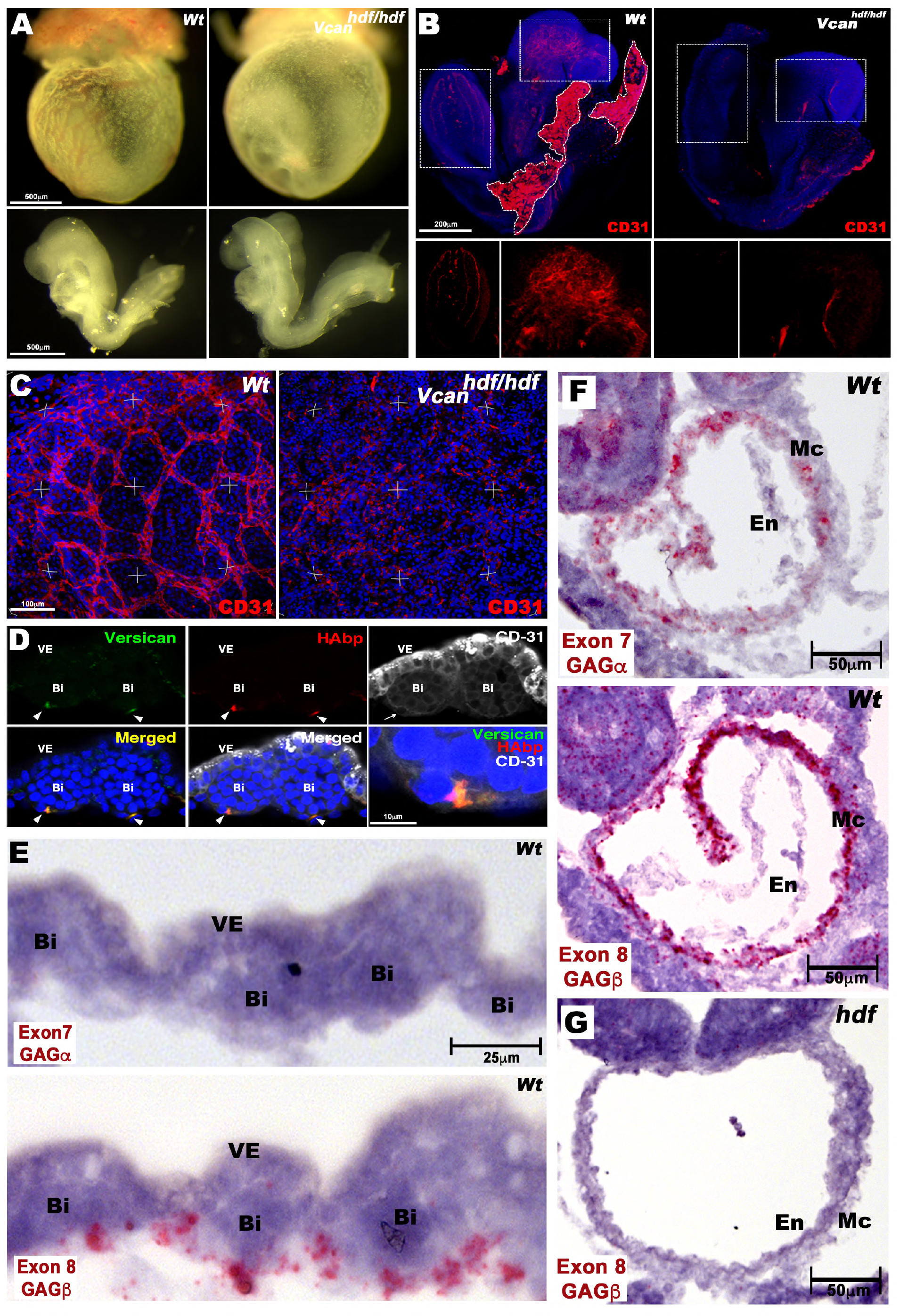
Impaired vasculogenesis in the *Vcan^hdf/hdf^* embryo and yolk sac. **(A)** E8.5 *Vcan^hdf/hdf^* yolk sacs are avascular yet dissected embryos appear morphologically similar to wild type. **(B)** Maximum intensity projections of whole mount E8.5 wild type and *Vcan^hdf/hdf^* embryos stained with anti-CD31 (red) and DAPI (blue). Boxed areas are shown at higher magnification with red channel only in the lower panels. Residual wild-type yolk sac is marked by a white dotted line (n=3 embryos from each genotype). **(C)** Three-dimensional (3D) maximum intensity projections of yolk sacs stained *en face* with anti-CD31 (red) show a well-formed vascular plexus in wild type yolk sac and random CD31+ cells in *Vcan^hdf/hdf^* yolk sac (n=3 yolk sacs from each genotype). **(D)** Cross section of E8.5 wild type yolk sac blood islands co-stained with versican (green), HAbp (red) and CD31 (white). Versican and HA co-localize with CD31+ cells on the mesothelial aspect of blood islands (arrowheads). The blood island imaged on the left is enlarged in the bottom right-hand panel (n=3 wild type yolk sacs). VE, visceral endoderm, Bi, blood island, **(E)** RNAscope in situ hybridization of E8.5 wild-type yolk sac shows expression of *Vcan* isoforms containing exon 8 (V1), but not exon 7 (V0, V2) in mesoderm adjacent to blood islands (Bi), VE, visceral endoderm. **(F)** *Vcan* exon 7 and exon 8 probes both hybridize to myocardium (Mc) of E8.5 wild-type embryos. En, Endocardium. **(G)** *Vcan* probes (exon 8 shown) do not hybridize to *Vcan^hdf/hdf^* heart. Scale bar in D=10μm, E= 25μm, 50μm in F-G.

### Versican and HA are associated with yolk sac Flk1^+^ cells

Flk1, a well characterized hematoendothelial progenitor marker indispensable for both vascular and blood lineage development [29], showed complete overlap with versican and HA on the mesothelial aspect of blood islands in wild-type E8.5 yolk sac (Fig. 3A). *Vcan*^hdf/hdf^ yolk sacs, in contrast, showed small blood islands with dramatically attenuated HA and Flk1 staining (Fig. 3A). *En face* confocal microscopy of whole-mount wild-type E8.5 yolk sacs revealed discrete patches of versican throughout the mesoderm (Fig. 3B). CD41, which is expressed upon specific commitment to the blood lineage [30], was strongly expressed in blood islands in the mid-mesoderm plane in wild-type yolk sacs (Fig. 3B). Versican did not precisely overlap with CD41+ cells and was restricted to intense patches corresponding to individual endothelial cells in wild-type yolk sacs, evident both in the surface (mesothelial) plane and cross-sections, whereas *Vcan*^hdf/hdf^ yolk sacs lacked CD41-stained blood islands entirely (Fig. 3B). High magnification images showed a few cells with weak CD41 staining in *Vcan*^hdf/hdf^ yolk sac mesoderm (arrowheads in Fig. 3B), contrasting with well-demarcated CD41+ wild-type blood islands. Combined, these results show that versican and HA distribution at E8.5 is restricted and specifically associated with uncommitted Flk1^+^ hemogenic endothelial cells, which are crucial for both blood and vascular development in the mouse embryo. Versican and HA are not specifically associated with subsequently differentiated blood cells (Fig. 3B,C).

**Figure 3.**
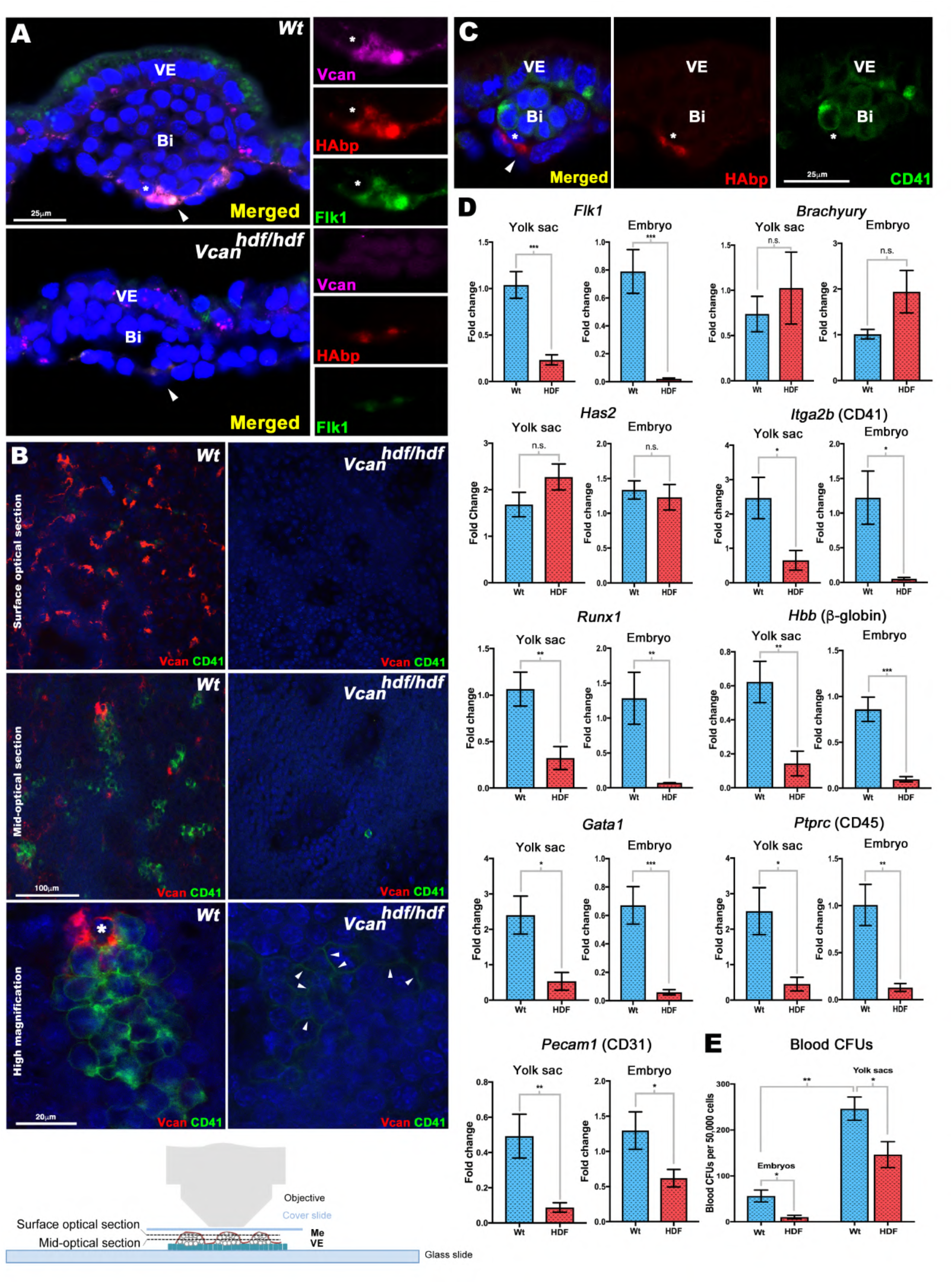
Versican and HA localize to Flk1+ cells and are essential for yolk sac blood island formation. **(A)** E8.5 wild type and *Vcan^hdf/hdf^* yolk sac cross-sections co-stained with versican (magenta), HAbp (red) and Flk1 (green). In wild type yolk sac versican-HA staining colocalizes with Flk1. In *Vcan^hdf/hdf^* yolk sac, blood islands (Bi) are smaller and both HAbp staining and Flk1 staining are weak. Arrowhead: versican-HA-Flk1 co-stained patches, asterisks: cell shown in high magnification in the right-hand panels (n=4 yolk sacs from each genotype). VE, visceral endoderm. **(B)** *En face* confocal imaging of E8.5 yolk sac versican (red) and CD41 (green) staining with the mesothelial aspect facing the objective (bottom). Surface optical sections show versican-rich loci throughout wild type yolk sac but absent in *Vcan^hdf/hdf^* yolk sac. The mid-optical image shows versican and CD41 co-stained cells in blood islands. Versican is associated with wild-type blood islands but does not overlap with CD41. No CD41+ cells were observed in *Vcan^hdf/hdf^* yolk sac. The high magnification image shows the distinct cell populations marked by versican and CD41. Arrowheads mark weak CD41 staining in *Vcan^hdf/hdf^* images (n=3 yolk sacs from each genotype). **(C)** Cross-section of an E8.5 wild type blood showing no overlap of HA (red) and CD41 (green) (n=3 yolk sacs). **(D)** qRT-PCR analysis of wild type and *Vcan^hdf/hdf^* yolk sacs and embryos shows significantly lower *Flk1* expression but not *Has2* or *Brachyury* expression in *Vcan^hdf/hdf^* mutants. CD41 *(Itga2b)* and *Runx1* expression were significantly lower in *Vcan^hdf/hdf^* yolk sac and embryos. Blood markers *β-globin*, *Gata1* and CD45 (*Ptprc*) and the vascular endothelial marker CD31 (*Pecam1*), were reduced in *Vcan^hdf/hdf^* yolk sacs and embryos (n=3 yolk sacs and embryos from each genotype, error bars= S.E.M;*, p<0.05; **, p<0.01; ***, p<0.001). **E.** Methylcellulose assay shows significantly fewer blood colony forming units (CFUs) in *Vcan^hdf/hdf^* yolk sacs and embryos (n=3 yolk sacs and three embryos from each genotype, error bars= S.D.,*, p<0.05; **, p<0.001).

### Versican is required for blood formation

qRT-PCR revealed significantly lower *Flk1* expression in *Vcan*^hdf/hdf^ yolk sac and embryos (Fig. 3D), in agreement with reduced Flk1 staining observed in the mutants (Fig. 3A). Neither expression of *Brachyury*, a mesoderm marker, nor *Has2*, encoding the major HA synthase in the embryo, were altered in *Vcan*^hdf/hdf^ yolk sacs or embryos (Fig. 3D). Expression of genes encoding blood lineage commitment markers *Itga2b* (CD41) and *Runx1* were similarly reduced in the *Vcan*^hdf/hdf^ yolk sacs and embryos, as well as erythroid and myeloid transcripts *Hbb* (*β*-globin), *Gata1* and *Ptprc* (CD45) (Fig. 3D). CD31 (*Pecam1*) mRNA expression was greatly reduced in the *Vcan*^hdf/hdf^ yolk sacs and embryos (Fig. 3D), consistent with the observed lack of vasculature (Fig. 2B). Methylcellulose colony formation assays demonstrated fewer colony-forming units (CFUs) in E8.5 *Vcan*^hdf/hdf^ embryos and yolk sacs (Fig. 3E), in agreement with fewer CD41-stained blood islands observed in the mutant yolk sacs.

### Versican and HA are associated with the earliest Flk1^+^ cells at gastrulation

Since Flk1^+^ cells arise at gastrulation, we analyzed versican and HA distribution in wild-type E7.5 mouse embryos. Versican and HA co-localized to the extraembryonic mesoderm in the blood island ring (Fig. 4 A,B), and colocalized with Flk1^+^ cells (Fig. 4 C,D). Correspondingly, RNA *in situ* hybridization of serial sections from several embryos showed consistent overlap between *Vcan*, *Has2* and *Flk1* expressing cells in the extraembryonic mesoderm (Fig. 4E, Fig.4 Supplement 1). *Vcan* exon 7-containing probes gave a weaker signal than exon 8 probes. In contrast, *Runx1*, which identifies blood lineage-committed cells, was expressed by cells in the extraembryonic mesoderm distinct from those expressing *Flk1*, *Vcan* and *Has2* (Fig. 4E, Fig.4 Supplement 1). Thus, versican-HA ECM is intimately associated with the hematoendothelial lineage emerging at gastrulation.

**Figure 4.**
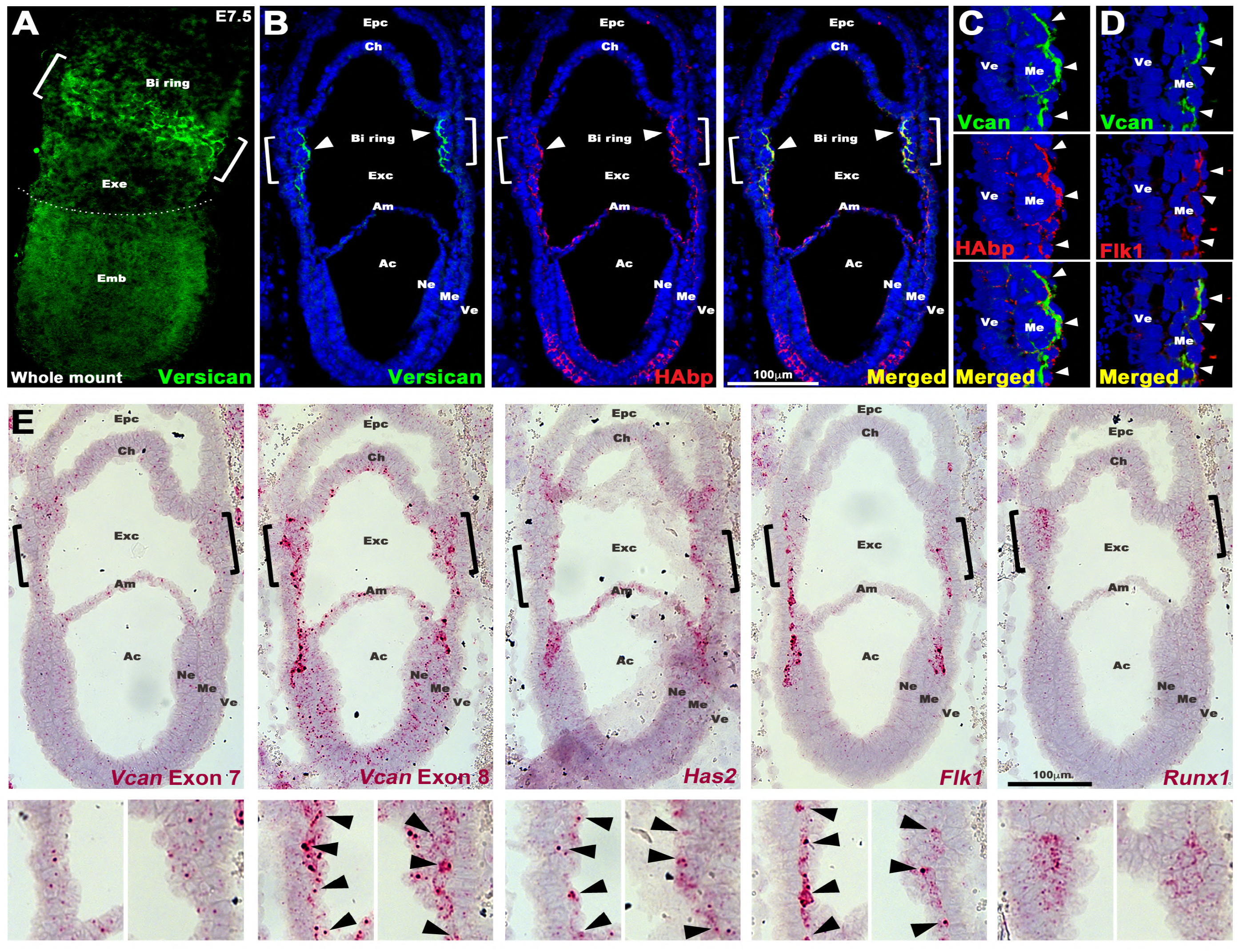
Versican and HA co-localize with Flk1+ cells from their origin at gastrulation. **(A)** Maximum intensity projection image of an E7.5 embryo shows strong versican staining (green) in the putative blood island (BI) ring (white brackets) in the proximal extraembryonic (Exe) region. Emb, embryo **(B-E)** Serial sections of E7.5 wild type embryos analyzed by immunostaining **(B-D)**, or by RNA in situ hybridization **(E)**. **(B-C)** Versican (green) and HAbp (red) colocalize in extraembryonic mesoderm corresponding to the blood island ring (white brackets and arrowheads). The blood islands on the left are shown at high magnification in **(C-D)**. Arrowheads show colocalization of versican, HAbp and Flk1 in extraembryonic mesoderm (N=4 embryos). Epc, ectoplacental cavity; Ch, chorion; Exc, exocelomic cavity; Am, amnion; Ac, amnion cavity; Ne, neural ectoderm; Me, mesoderm; Ve, visceral endoderm. **(E**) *In situ* hybridization for *Vcan* exon 7, *Vcan* exon 8, *Has2*, *Flk1* and *Runx1*. *Vcan* exon 8, *Has2* and *Flk1* mRNAs have near-identical expression patterns (red) corresponding to primitive streak cells migrating toward extra-embryonic mesoderm. *Vcan* exon 7 (GAG*α*) shows weaker overlapping expression. *Runx1* marks committed blood cells (n=4 embryos). Scale bars in **B,E**= 100μm.

Data mining of a single cell RNA sequencing (scRNA-seq) atlas of early (6.5 to 8.5 day-old) mouse embryos [31] supported specific *Vcan* expression by hematoendothelial progenitors. High *Vcan* expression was first detected in hematoendothelial progenitors emerging at E6.75, as well as in nascent and uncommitted mesoderm (Fig. 4 Supplement 2A-C). *Vcan* was expressed in *Kdr* (encoding *Flk1*)-expressing cells at E6.75, E7.0 and E7.5, spanning mouse gastrulation (Fig. 4 Supplement 2A-C). *Kdr*^+^ cells also expressed *Has2* and the gene encoding another HA receptor, CD44 (Fig. 4 Supplement 2B). These observations suggest a potential cell autonomous role for versican in Flk1^+^ cells, bound to HA and thus to the cell surface via CD44 and HAS2. After E7.5 (e.g., at E 8.5), *Vcan* was broadly expressed in mesoderm, the developing brain, allantois and cardcardiomyocytes (Fig. 4 Supplement 2C) and *Kdr* expression extended into endothelial cells, although hematoendothelial progenitors subsequently continued to express *Vcan* and *Kdr* (Fig. 4 Supplement 2C). scRNA-seq data showed that HA link proteins (*Hapln1-4*) were not expressed in hematoendothelial progenitors (Fig. 4 Supplement 3A), and the major versican-binding link protein, HAPLN1, was absent in E7.5 embryo sections (Fig. 4 Supplement 3B). Staining for ADAMTS-cleaved versican (using anti DPEAAE) revealed no staining in wild-type blood islands, suggesting that versican does not undergo proteolysis by ADAMTS proteases at this time (Fig.4 Supplement 3C).

### Loss of versican dramatically reduces yolk sac and embryo HA levels

In addition to reduced HA staining in *Vcan*^hdf/hdf^ yolk sac blood islands, we observed dramatic loss of HAbp staining throughout *Vcan*^hdf/hdf^ embryos (Fig. 5A). Since *Has2* mRNA expression was unaltered in *Vcan*^hdf/hdf^ embryos and yolk sacs (Fig. 3D), reduced HABP staining may have resulted from increased breakdown in situ, or alternatively, extraction of unliganded HA from tissue sections during staining. Therefore, fluorophore-assisted carbohydrate electrophoresis (FACE) [32–34] was undertaken on snap-frozen whole embryos (Fig. 5B). FACE analysis demonstrated that versican is a major CSPG in E8.5 embryos, contributing nearly all unsulfated (0S) CS, since both the overall level of CS and its 0S form were dramatically reduced in *Vcan*^hdf/hdf^ embryos (Fig. 5B). In addition, HA-FACE showed reduction of HA compared to wild-type littermates, supporting HA loss intrinsically within the embryo rather than during sample preparation (Fig. 5B). Consistent with the role of versican and HA as an expansile complex, craniofacial mesenchyme and myocardium were severely compacted in *Vcan*^hdf/hdf^ embryos (Fig. 5C). We also asked whether the absence of versican affected fibronectin, a major embryonic ECM component that is essential for angiogenesis and observed a dramatic increase both in fibronectin staining and mRNA in *Vcan*^hdf/hdf^ embryos (Fig. 5D, Fig. 5 Supplement 1A,B), which is presently unexplained. In combination with unaltered *Has2* expression, these findings suggest increased HA turnover in *Vcan*^hdf/hdf^ embryos. TMEM2 was recently identified as a major extracellular hyaluronidase [35–37] and qRT-PCR revealed significantly raised*Tmem2* expression in *Vcan*^hdf/hdf^ yolk sac and embryos (Fig. 5E). indicating that hyaluronan catabolism was activated in the absence of versican.

**Figure 5.**
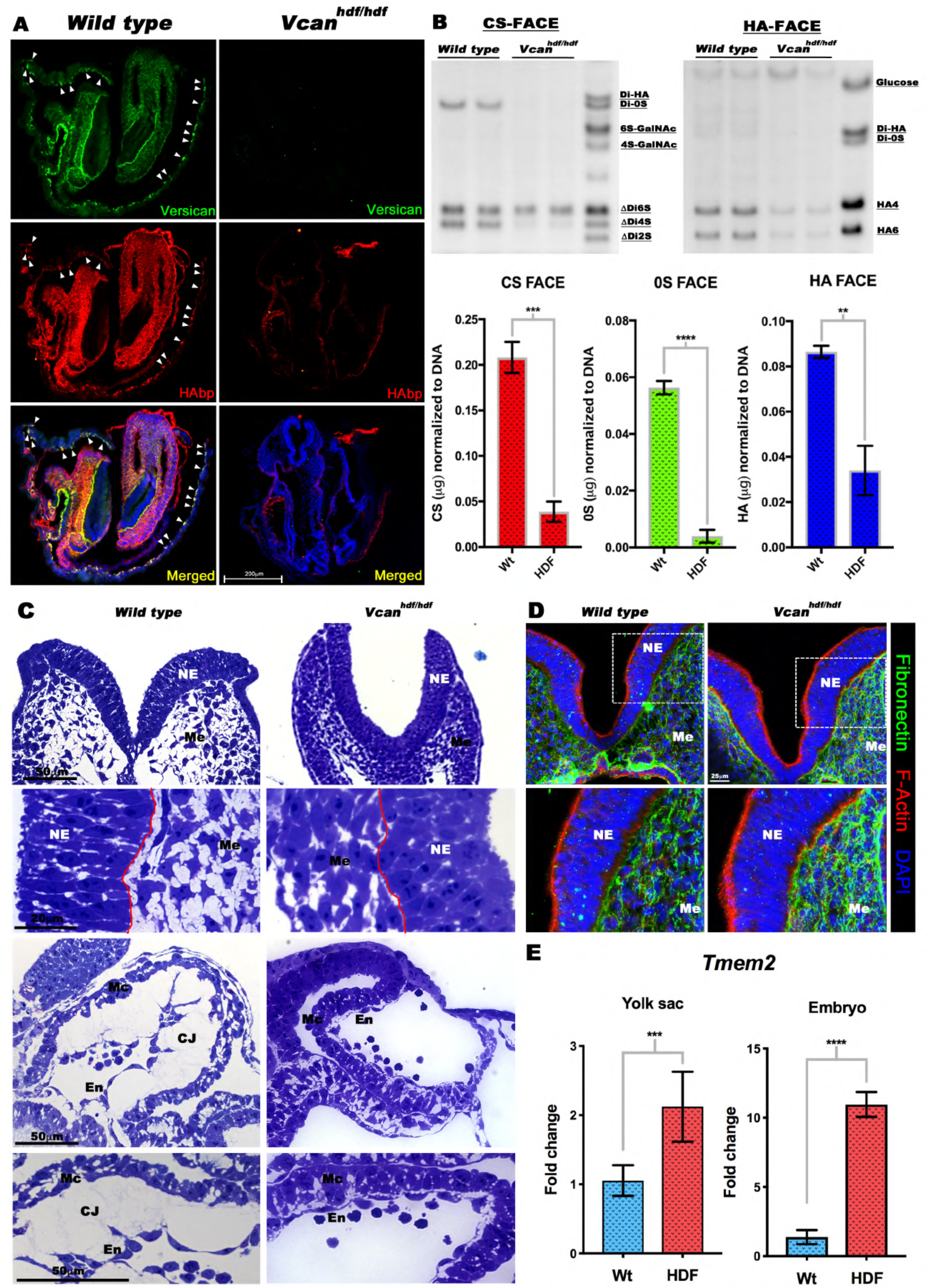
*Vcan* inactivation results in loss of hyaluronan. **(A)** Versican (green) and HAbp (red) staining of E8.5 wild type and *Vcan^hdf/hdf^* embryos showing severely reduced HAbp staining in *Vcan^hdf/hdf^* embryos. Arrowheads indicate versican and HAbp-stained yolk sac blood islands. **(B)** *(Top)* Fluorophore-assisted carbohydrate electrophoresis (FACE) analysis for chondroitin sulfate (CS-FACE) and hyaluronan (HA-FACE) in E8.5 littermate embryos. Versican is the major CS-proteoglycan in E8.5 embryos. HA is significantly reduced in *Vcan^hdf/hdf^* embryos. *(Bottom)* Quantification of CS and HA FACE, normalized to DNA (n=4 embryos from each genotype, error bars= S.E.M.,**, p<0.01; ***, p<0.001; ****, p<0.0001). **(C)** 1 μm thick, toluidine blue-stained Eponate 12 sections from E8.5 wild type and *Vcan^hdf/hdf^* embryos show compaction of craniofacial mesenchymal cells (Me) and loss of cardiac jelly (CJ) between the myocardium (Mc) and endocardium (En). The red line indicates the boundary of neural epithelium (NE) with mesenchyme. **(D)** E8.5 wild *Vcan^hdf/hdf^* embryo sections stained with F-actin (red) and fibronectin (green) show stronger fibronectin staining. **(E)** qRT-PCR shows increased *Tmem2* transcription in *Vcan^hdf/hdf^* embryos and yolk sacs (N=3 embryos and yolk sacs from each genotype, error bars= S.E.M.;***, p<0.001; ****, p<0.0001). Scale bar in **A**= 200μm, 50μm and 20μm in **C**.

### HA deficiency impairs vasculogenesis

Since HA was lost in the absence of versican, we inquired if the converse were true, and whether HA-deficiency indeed led to defective yolk sac vasculogenesis, as previously suggested by morphology of *Has2* germline mutants [26]. *Has2*^−/−^ embryos die by E 9.5 due to heart defects [26] whereas *Has1^−/−^* and *Has3^−/−^* mice are both viable and do not exhibit noticeable embryonic phenotypes [38, 39]. To completely deplete HA from developing embryos, we generated triple compound embryos lacking all three *Has* genes (*Has1^−/−^;Has2^−/−^;Has3^−/−^*; referred to as *Has1-3^TKO^*). We generated HA-deficient mouse embryos by intercrossing *Has1+3^−/-^; Has2^+/-^* mice. At E9.5, *Has1-3^TKO^* embryos, like *Vcan*^hdf/hdf^ and *Has2*-epiblast deleted embryos [26] showed loss of yolk sac vasculature (Fig. 6A). HA staining in the *Has1-3^TKO^* yolk sac and embryos was absent, as expected and the sections also showed a near-complete lack of versican staining (Fig. 6B,C). In contrast, immunostaining for cleaved versican (anti-DPEAAE) showed robust and widespread versican processing in the absence of HA (Fig. 6 B,D). Epiblast-specific *Has2* conditional deletion (*Sox2*Cre;*Has2^Fl/Fl^*) showed a similar loss of yolk sac vasculature and cardiac defects (Fig.6 Supplement 1A). RNA in situ hybridization showed a low level *Has2* expression in these embryos and slightly reduced *Vcan* transcript levels (Fig. 6 Supplement 1B). Phenocopying *Has1-3^TKO^* embryos, the *Sox2*Cre; *Has2^Fl/Fl^* embryos and yolk sacs also showed loss of both HA and versican staining and increased ADAMTS-mediated versican cleavage (Fig. 6 Supplement 1C). Taken together, these data suggest that HA may protect versican from ADAMTS-mediated degradation.

**Figure 6.**
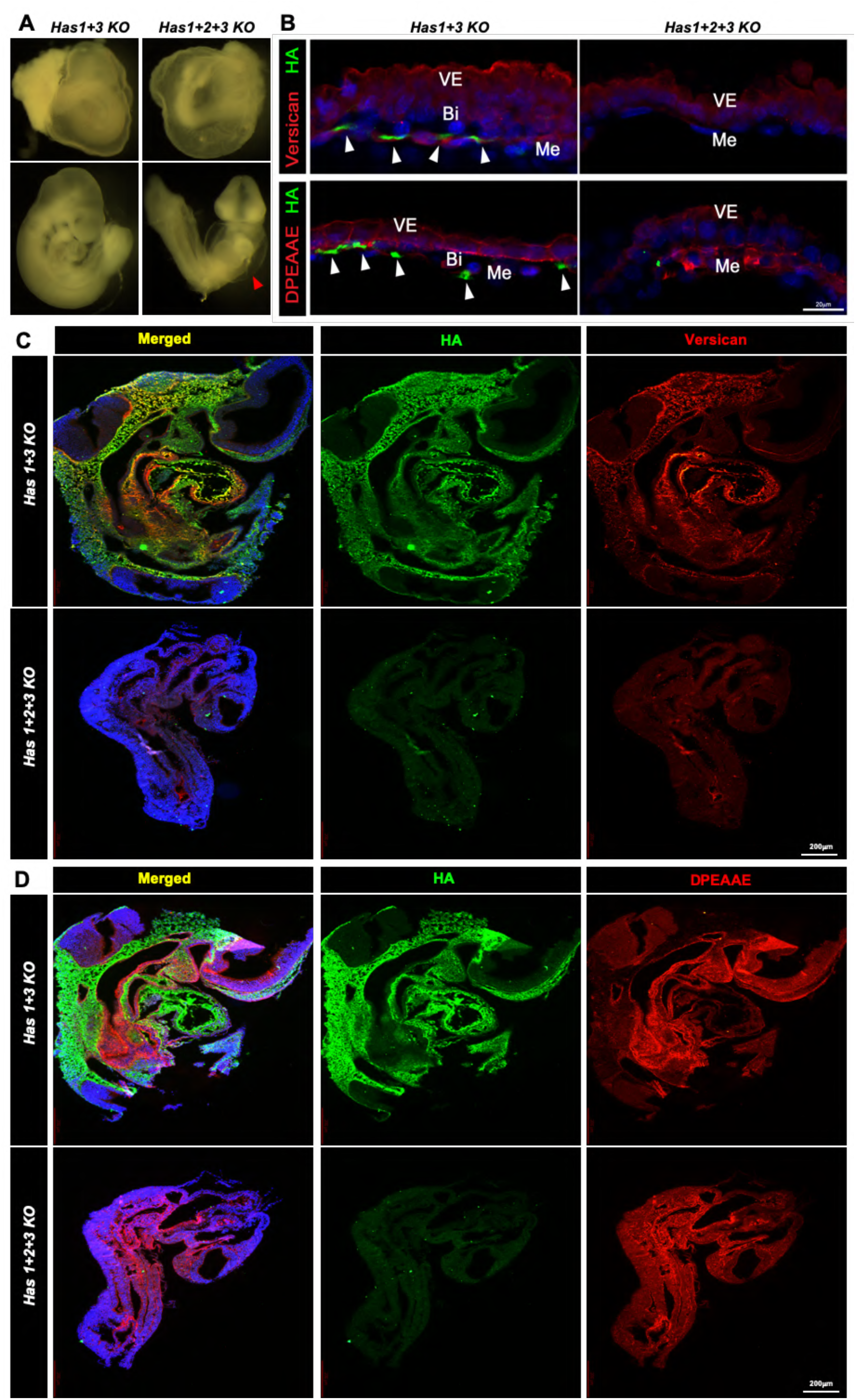
Versican is severely reduced in *Has1+2+3* null embryos. **(A)** E9.5 *Has1+3* null embryos (control) and *Has1+2+3* null embryos showing the dilated pericardial sac in the latter (arrowhead) similar to *Vcan^hdf/hdf^*. **(B)** *Has1+3* null yolk sac (control) and *Has1+2+3* null yolk sac co-stained with HAbp (green) and versican or cleaved versican (DPEAAE) (red) show loss of both HA and versican although cleaved versican (DPEAAE, red, bottom) is present in *Has1+2+3* null yolk sac. **(C-D)** *Has1+3* null yolk sac (control) and *Has1+2+3* null embryos co-stained with HAbp and versican **(C)** or cleaved versican (DPEAAE) **(D)**, show loss of HA and versican staining and weaker DPEAAE staining in the *Has1+2+3* null embryos. Scale bars = 200μm in in **C**,**D**.

### Embryoid bodies formed by *Vcan*-null ES cells have impaired angiogenesis and hematopoiesis

Since the dramatic loss of vasculature could result from perturbed hemodynamics due to cardiac defects in *Vcan*^hdf/hdf^ embryos, we undertook an orthogonal (in vitro) approach for evaluation of versican in vasculogenesis and hematopoiesis. We used gene editing by CRISPR/Cas9 [40, 41] to introduce *Vcan-*null mutations in R1 mouse embryonic stem cells (mESC) [42]. *Vcan* exon 2 (containing the start codon) and exon 3 (start of the G1 domain) were targeted independently to obtain mESC clones D8 and F9 respectively, each with defined frameshift mutations that generated null alleles (Fig. 7A). Since exons 2 and 3 are included in all *Vcan* splice isoforms, no versican was produced, as shown by western blot with anti-GAG*β* antibody (Fig. 7B). The mutated clones retained normal expression of *Oct4, Sox2, Nanog* and *C-myc* (Fig. 7C) indicating unaltered pluripotency, validating their suitability for *in vitro* differentiation. *Vcan*-null and wild-type ES cells were allowed to form embryoid bodies (EBs), in which random differentiation into various lineages occurs. RT-qPCR analysis of *Has2* showed no change in *Vcan-*null EBs (Fig. 7 Supplement 1A) while the mesoderm differentiation marker *Brachyury/T* showed a significant decrease in the F9 but not the D8 *Vcan-*null EBs (Fig. 7 Supplement 1B). Similar to *Vcan*^hdf/hdf^ embryos, *Vcan* null embryoid bodies showed reduced *Kdr* expression (Fig. 7 Supplement 1C), and the blood lineage commitment markers *Itga2b* (CD41) and *Runx1* (Fig. 7 Supplement 1D), as well as of differentiated-blood and vascular endothelial markers (Fig. 7 Supplement 1E, F). Since EB differentiation is random and independent of a closed pulsatile circulation, the data from embryoid bodies and the mutant mice together suggests that versican acts directly on Flk1^+^ hematoendothelial progenitors rather than indirectly *via* abnormal cardiac development.

**Figure 7.**
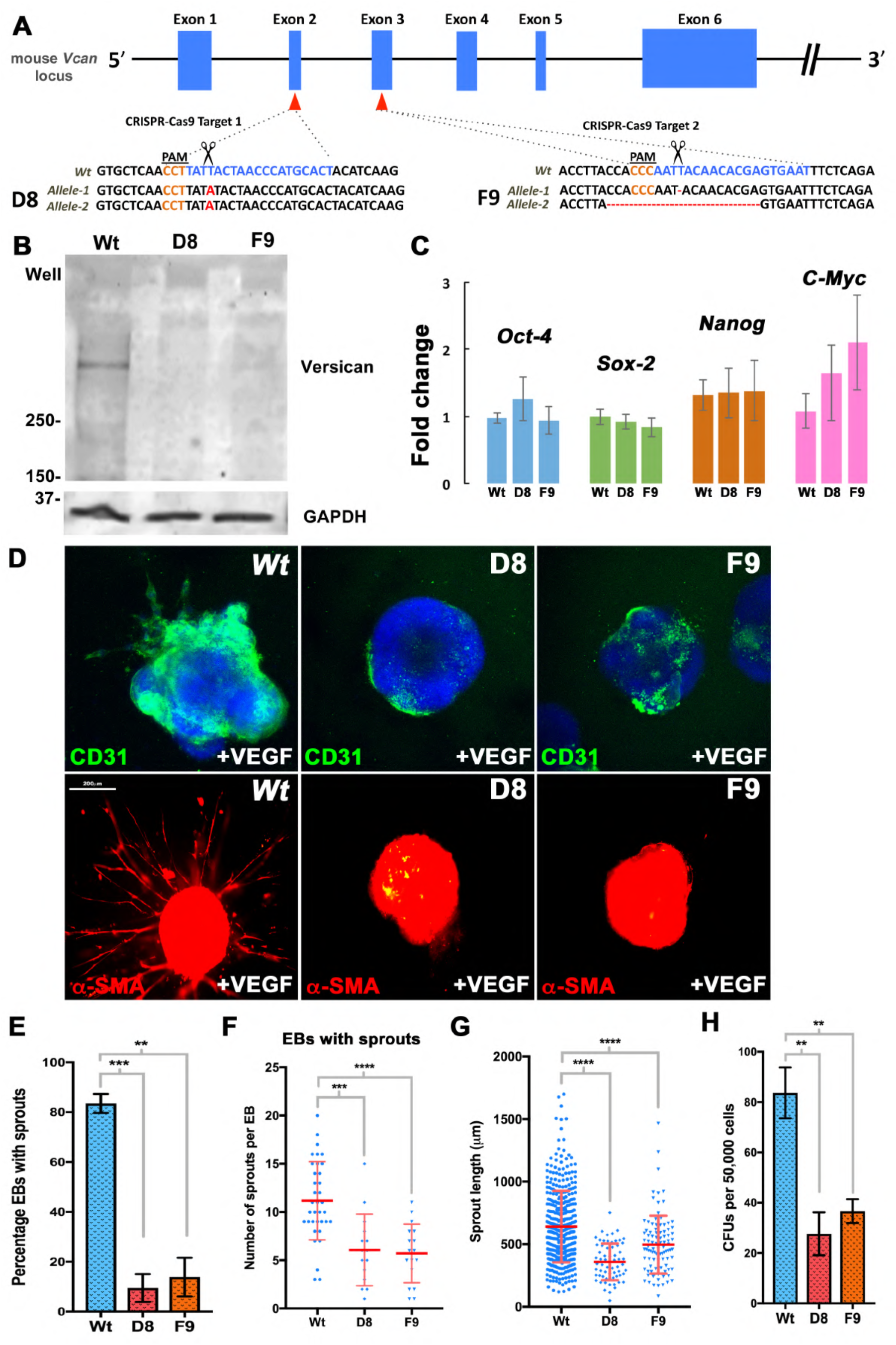
Embryoid bodies (EBs) from *Vcan*-null mouse embryonic stem cells (mESCs) are unresponsive to VEGF and form few blood colonies. **(A)** *Vcan* locus showing targeting by independent guide RNAs (gRNAs) targeting exons 2 and 3. Scissors indicate the Cas9 cleavage site 3 bp from the protospacer adjacent motif (PAM, orange lettering). An exon 2 mutant mESC clone (D8) had a homozygous 1 bp insertion (bold red letters) and an exon 3 mutant clone (F9) had heterozygous targeting: deletion of T/A in one allele and a 19-bp deletion in the other resulting in frame-shifts. **(B)** Western blot of 10-day differentiated EBs shows no versican in EBs derived from clones D8 and F9. **(C)** qRT-PCR analysis showing unaffected expression of pluripotency markers *Oct4*, *Sox2*, *Nanog* and *C-myc* in D8 and F9 (n=3 independent batches of EBs from each genotype, error bars= S.E.). **(D)** 4-day-old *Vcan*-null embryoid bodies embedded in collagen I and treated with VEGF_165_ for 12 days lack vascular sprouting identified by CD31 (green) or *α*-SMA (red) immunostaining (n=3 independent batches of EBs from each genotype). **(E)** D8 and F9 EBs show significantly fewer vascular sprouts (n=3 independent batches of EBs from each genotype, error bars= S.E. **, p<0.01; ***, p<0.001). **(F)** Fewer sprouts/EB were seen in D8 and F9 EBs. (n=3 independent batches of EBs from each genotype, error bars= S.D. ***, p<0.001; ****, p<0.0001). **(G)** Significantly shorter sprouts in *Vcan* null lines (n=3 independent batches of EBs from each genotype, error bars= S.D. ****, p<0.0001). **(H)** Methylcellulose colony assay using dissociated 10-day old EBs shows significantly fewer colonies in *Vcan* null EBs. (N=3 independent batches of EBs from each genotype, error bars=S.D. **, p<0.01). Scale bar in D is 200 μm.

Additionally, we separately evaluated the endothelial and hematopoietic potential of *Vcan*-null ESC. Sprouting angiogenesis was induced by culturing 4-day differentiated EBs in 3-dimensional collagen I gels and treatment with VEGF-A_165_ for 12 days [43]. Robust angiogenic sprouting identified by CD31 and smooth muscle *α*-actin staining was seen in wild-type EBs, but not in the majority of *Vcan-*null EBs (Fig. 7 D,E). The few *Vcan-*null EBs with sprouting had fewer sprouts per EB, and these were significantly shorter than wild-type sprouts (Fig. 7 F,G). When 10-day old EBs were disaggregated and the cells were plated in a 3-dimensional methylcellulose matrix containing a complete set of blood differentiation cytokines, significantly fewer blood colonies were formed by *Vcan* null EBs (Fig. 7 H).

### The versican-HA complex sequesters growth factors essential for vasculogenesis and hematopoiesis

These observations raised questions about the underlying mechanisms by which versican-HA matrix would support survival and expansion of Flk1^+^ cells. An *ADAMTS9* null RPE1 cell line (D12) lacks ADAMTS9, a key versican-degrading protease [22] and thus demonstrates constitutively strong versican staining in ECM (Figure 8A). HABP staining of these cells demonstrated stronger HA staining than in wild type RPE1 cells, with formation of long HA cables decorated with versican (Fig. 8A, Fig. 8 Supplement 1A,B); such cables are well-known to occur in 25 mM glucose-containing medium [44] [45]. qRT-PCR revealed *HAS2* & HAS3 expression were reduced in D12 vs. wt (Fig. 8B) with no change in *HAS2* expression but drastic reduction of *TMEM2* expression was observed in the RPE1-D12 cells (Fig. 8B). Intriguingly, upon siRNA-mediated versican depletion [46, 47], RPE1-D12 cells showed dramatically reduced versican and HA staining and absence of HA/versican cables (Fig. 8C, Fig. 8 Supplement 2A). Conversely, qRT-PCR showed increased *TMEM2* expression after *VCAN* knockdown (Fig. 8 Supplement 2B), suggesting regulation of the HA-depolymerase *TMEM2* by the level of versican as a possible mechanism of concurrent HA loss.

**Figure 8.**
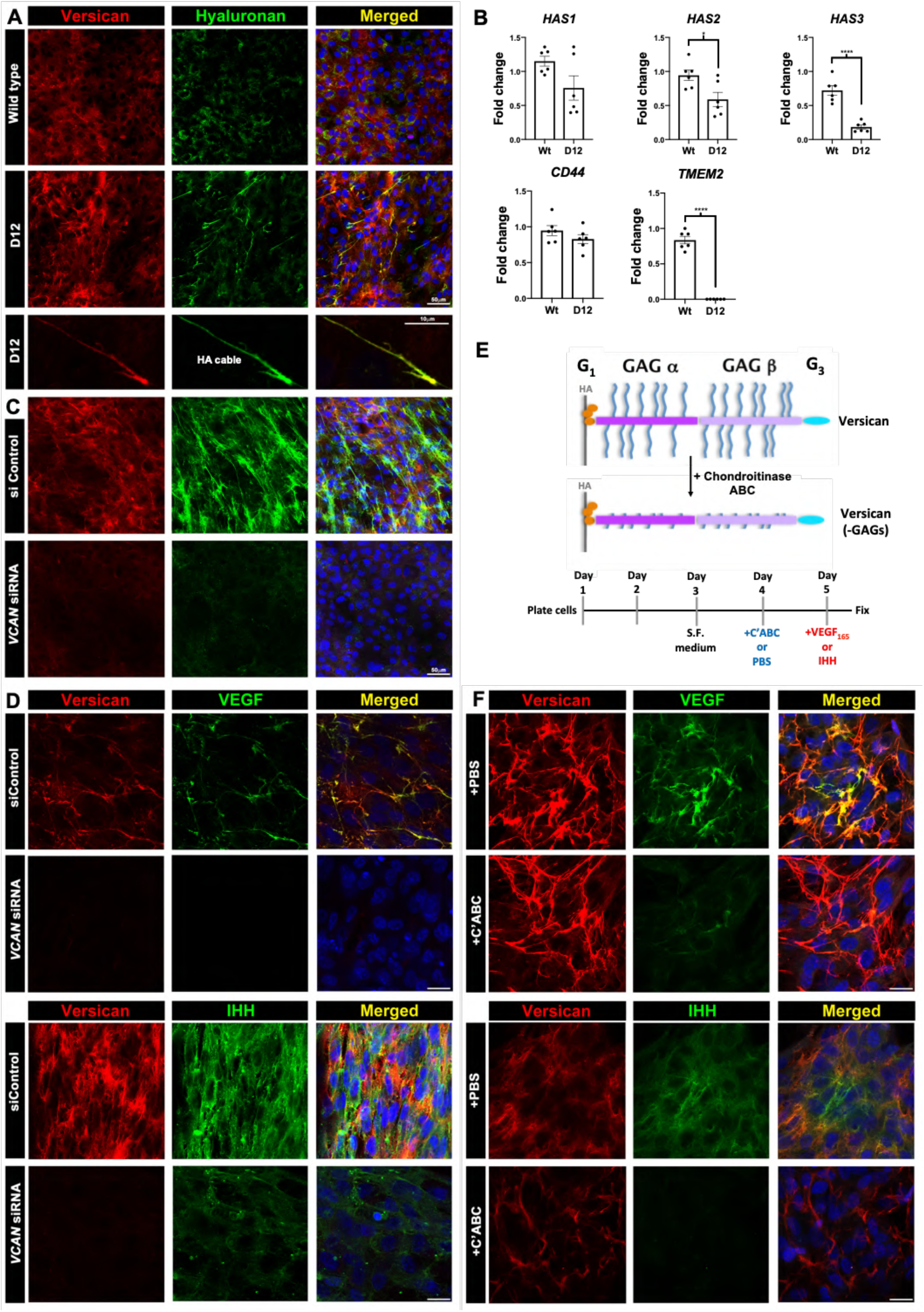
Versican regulates HA abundance and cable formation and sequesters VEGf and Ihh *via* chondroitin sulfate (CS) chains in a cell culture model. **(A)** Increased versican (red) and HAbp staining (green) in *ADAMTS9*-deficient (D12) RPE-1 cells. Versican and HA co-stained cables are formed in D12 cells. **(B)** qRT-PCR showing no change in *HAS1-3* or *CD44* but dramatically reduced *TMEM2* expression in D12 cells. (n=3 independent RNA extractions, error bars= S.E.M., *, p<0.05; ****, p<0.0001). **(C)** *VCAN* knockdown reduces both versican (red) and HA staining in D12 cells. **(D)** Recombinant VEGF_165_ or Ihh (green) co-staining with versican (red) in control siRNA-transfected D12 cells is lost in *VCAN* siRNA-transfected D12 cells. **(E)** The schematic illustrates chondroitinase ABC removal of versican GAG chains and the experimental timeline used. **(F)** Fluorescence microscope of VEGF and Ihh with versican, with or without chondroitinase ABC treatment prior to addition of recombinant VEGF_165_ or Ihh (green) shows reduced binding of both growth factors after CS-chain removal, but does not affect versican core protein staining (red).

Next, to test if versican can sequester growth factors known to regulate vasculogenesis, we treated wild-type and RPE1-D12 cells with increasing concentrations of recombinant VEGF_165_ and Ihh. Each showed dose-dependent punctate staining in RPE1-D12 monolayers which co-localized with the versican-HA cables, imaged by super-resolution fluorescent microscopy. (Fig. 8D, Fig. 8 Supplement 3A-B, Fig. 8 Supplement 4A-B). Upon *VCAN* depletion in RPE1-D12 cells both VEGF_165_ and Ihh staining were significantly decreased (Fig. 8D). Enzymatic removal of proteoglycan chondroitin sulfate chains prior to addition of recombinant VEGF_165_ and Ihh to D12 cultures also dramatically reduced VEGF_165_ and Ihh staining intensity without affecting versican core protein staining (Fig. 8E,F), suggesting that versican CS chains mediated VEGF and Ihh sequestration.

## Discussion

Although versican is widely expressed during organogenesis and in adult tissues [14], the chronology of its expression in early vascular development undertaken here and independently derived from recently published single cell RNA-seq data, shows specific association with Flk1^+^ cells from their earliest origin in the gastrulating embryo until establishment of yolk sac vasculature and primitive hematopoiesis. Analysis of *Vcan* mutant mice *in vivo* and *Vcan-*null embryoid bodies *in vitro* suggest that versican has an indispensable role in vasculogenesis and primitive hematopoiesis. Taken together, RNA *in situ* hybridization data, scRNA-seq analysis of Flk1^+^ cells and immunostaining of embryos suggests that versican is a product of Flk1^+^ cells that associates with them via HA and CD44 to form a crucial, possibly cell-autonomously acting ECM. Furthermore, in vitro analysis undertaken in REP1-D12 cells for their high versican levels, showed dose-dependent binding of VEGF and Ihh to the CS-chains of versican.

Two novel, unexpected findings of this study were the substantial loss of embryonic and extraembryonic HA in the absence of versican and reciprocally, of versican in the absence of HA. Also unexpected was the transcriptional effect of versican expression on TMEM2. As in *Vcan*^hdf/hdf^ yolk sac, avascularity was previously noted in *Has2* null yolk sac [26], although it was not studied further. *Has2* and *Vcan* null cardiac defects are similar, which suggested that an obligate versican-HA complex in cardiac jelly and endocardial cushions was required for cardiac morphogenesis [24–26]. The present work similarly demonstrates that a versican-HA complex, rather than versican alone, is necessary for vasculogenesis and primitive hematopoiesis (Fig. S12). HA-versican aggregates have a net negative charge, and exert swelling pressure *via* absorption of water (the Gibbs-Donnan effect) [48]. As shown here and previously, embryonic tissues compact in their absence [24, 49]. Although versican-deficient yolk sac showed extensive structural disorganization at E9.5, it is clear that the association and effect of versican vis-à-vis Flk1^+^ cells occurs much earlier, i.e., shortly after gastrulation. Reduced vasculogenesis, angiogenesis and hematopoiesis in *Vcan-*null embryoid bodies supports a direct, local role of versican on hematoendothelial progenitors than a secondary effect of cardiac anomalies on vasculogenesis, or of yolk sac disorganization on blood island formation. We therefore conclude that versican-HA aggregates act directly and as early as E6.75 within the microenvironment of Flk1^+^ cells.

HA, which is extruded directly from HA synthases on the plasma membrane, is decorated with versican, and is retained at the cell surface via HA synthases or CD44 and other HA receptors [50]. *Cd44* is coordinately expressed with *Vcan* and *Has2* by Flk1^+^ cells during early embryogenesis. Nevertheless, *Cd44* null mice survive and are not known to have vasculogenesis or primitive erythropoiesis defects [51], suggesting that HAS2 acting as a *de facto* receptor, may participate in forming a cell-associated versican-HA complex. The versican-HA pericellular matrix has a well-established impact on different cells. We previously found that the amount of versican in the pericellular matrix in fibroblasts and vascular and myometrial smooth muscle cells [9, 21, 52] was a determinant of phenotype modulation. Versican is anti-adhesive [9, 52], which may allow Flk1^+^ cells to migrate and proliferate efficiently after their emergence at gastrulation. Another possible role of versican-HA is sequestration of vasculogenic factors such as VEGF-A. Versican binding to VEGF-A and Ihh *via* its CS-chains (this study) or G3 domain [19], may generate high concentrations around Flk1^+^ cells. Versican is associated with Flk1^+^CD41^−^ cells at their origin, but not with the Flk1^−^CD41^+^ yolk sac cells subsequently, suggesting that the versican-HA pericellular ECM specifically regulates the fate of Flk1^+^ CD41^−^ cells. We conclude that as the dominant CSPG in the embryo, specifically localized to the vicinity of Flk1^+^ cells, versican may sequester essential factors such as VEGF-A and Ihh to provide a high local concentration that sustains the Flk1^+^ population.

Although lack of versican led to higher *Fn1* expression, *Vcan*^hdf/hdf^ defects are unlikely to result from excess fibronectin, because previous work has shown that deficiency, not excess of fibronectin or the fibronectin receptor subunit *α*5 integrin results in reduced embryo and yolk sac vascularity [27, 53]. Since versican binds fibronectin through the G3 domain [19], we conclude that the three major components of the provisional embryonic ECM (versican, HA and fibronectin) are each crucial for establishment of the first blood vessels in the embryo. In the absence of versican, the HA catabolism rate *via* TMEM2 likely exceeds the HA synthesis rate since *Has2* mRNA levels were relatively unchanged. It was previously noted that HA deposition was reduced to 85% of normal in fibroblasts taken from a mouse hypomorphic *Vcan* mutant (*Vcan*^Δ3/Δ3^) having 75% reduction of versican [54]. In association with reduced HA deposition, *Vcan*^Δ3/Δ3^ fibroblasts had accelerated senescence, suggesting a possible mechanism for the lack of Flk1^+^ cells in *Vcan*^hdf/hdf^ yolk sac, although this study did not specifically address the fate of Flk1+ cells.

We observed stronger association of exon 8-containing *Vcan* transcripts than exon 7-containing transcripts with blood islands at gastrulation and exon 7 transcripts were absent at E8.5. Autosomal dominant splice site mutations affecting exon 8 (leading to its exclusion) and exon 8 deletions in humans cause Wagner syndrome [55–57], which is characterized by impaired vision and defects of the ocular vitreous and retina, but lacks consistent extra-ocular manifestations. Exon 7 inactivation in mice led to specific neural anomalies and subtle cardiac anomalies [58, 59]. Thus, neither individual exon mutation in mice or humans is associated with embryonic lethality, defective vasculogenesis or impaired hematopoiesis. We conclude that both exon 7 and exon 8-containing transcripts, possibly included in V0, the transcript containing both exons, are required for vasculogenesis and hematopoiesis.

The GAG*β*domain encoded by exon 8 has a unique N-terminal sequence, which is cleaved by ADAMTS proteases at the E^441^-A^442^ peptide bond in several contexts, notably cardiac valve development, interdigital web regression, umbilical cord development, neural tube and palate closure and myometrial activation [9, 22, 60]. The resulting N-terminal versican V1 fragment, G1-DPEAAE^441^, named versikine, is bioactive in interdigital web regression and myeloma growth [61, 62]. *Vcan* knock-in mouse mutants in which E^441^-A^442^ was mutated to render it uncleavable, complete gestation and havenormal yolk sac avascularity, suggesting that E^441^-A^442^ cleavage is not involved in hematoendothelial development ([63] and Nandadasa et al., manuscript in preparation).

With the added role in vasculogenesis and hematopoiesis elucidated here, it is not an exaggeration to state that versican is crucial for development of the entire circulatory system. Indeed, versican-HA matrix may also have a broad role in formation of vasculature by angiogenesis in a variety of physiological and disease settings. Recent work found that syngeneic B16F10 tumors in adult *Vcan^hdf/+^* mice had significantly impaired angiogenesis and reduced growth [64]. Relevant to the overlap of versican-HA with Flk1 expression, a recent study utilizing the same cancer model in *Flk1*+/- mice also found reduced angiogenesis during tumor growth [65]. Furthermore, VEGF binding of the versican-HA ECM which is quantitatively increased in the presence of high glucose, may be relevant to diabetic retinopathy, a common complication of diabetes where the increased activity of VEGF is well-established and indeed, a current target of treatment [66].

## Methods

### Mice

*Vcan*^Tg(Hoxa1)1Chm^ (*Vcan*^hdf^) mice [25] (Figure-1 Supplement 1) were obtained under a material transfer agreement from Roche. Generation of *Has1^−/−^;Has3^−/−^* double knockout mice was described previously [67]. A *Has2*-null allele (*Has2^−^*) was created from a *Has2^flox^* allele [68] by crossing *Has2^flox/flox^* mice with the germline deleter *Meox2-Cre* mice [69]. All of these mouse lines were backcrossed to C57BL/6J for more than 10 generations. *Has1^−/−^;Has2^+/–^;Has3^−/−^* mice were bred from these mutant mice. Triple *Has* knockout embryos were generated by crossing *Has1^−/−^;Has2^+/–^;Has3^−/−^* female and male mice. In a second approach to generate HA-deficient mouse embryos, *Has2^Fl/Fl^; Sox2Cre^tg^* mice were generated by crossing *Has2^Fl/Fl^* males with *Has2^+/Fl^; Sox2Cre^tg^* females. Mouse experiments were conducted with IACUC approval (Cleveland clinic protocols 2015:1530 and 2018:2045). Mice were maintained in a fixed light-dark cycle with food and water *ad libitum*. For genotyping E8.5 embryos with intact yolk sacs, the allantois was dissected out and lysed in 10 μl DirectPCR (Tail) digest reagent (Qiagen, catalog no. 102-T) supplemented with 1 μl of proteinase K overnight at 55°C. Tails were used to genotype E9.5 embryos. *Vcan^hdf^* and wild-types were identified with a specific genotyping strategy based on the genetic interruption (Figure-1 S1). Details of the *Has1-3* mutant mouse genotyping is provided in the Supplemental File. *Vcan^hdf/hdf^* embryos were compared to wild-type littermates in all experiments.

### Mouse embryonic stem cell (mESC) culture

R1 mESC [42] were cultured on 0.3% type B gelatin (Sigma-Aldrich, catalog no. G9382) coated 60 mm cell culture plates in Iscove’s Modified Dulbecco’s Medium (IMDM) containing 4mM L-glutamine and 1mM sodium pyruvate, supplemented with 20% fetal bovine serum (Hyclone, catalog no. SH30071), 0.1 mM cell culture grade 2-mercaptoethanol (Gibco, Life Technologies, catalog no. 21985), 0.1mM nonessential amino acids (Gibco, Life Technologies, catalog no. 11140-050), 50 μg/mL penicillin/streptomycin and 1×10^6^ units/mL leukemia inhibitory factor (LIF; ESGRO, EMD Millipore, catalog no. ESG1106) in a humidified 5% CO_2_, 37°C environment. mESC were maintained at 60-80% confluence, with daily medium change and passaged every other day in a 1:5 split.

### CRISPR/Cas9 targeting of mESC *Vcan*

2.5 μg of CRISPR/Cas9 plasmids in the U6gRNA-Cas9-2A-GFP vector, targeting *Vcan* exon 2 or exon 3 (Sigma-Aldrich, target IDs MM0000080027 and MM0000080028) were transfected into R1 mESCs at 60-80% confluence in 6-well plates coated with 0.3% gelatin, using FuGene 6 (Promega, catalog no. E2691). 24 hrs post-transfection, individual GFP+ mESCs were sorted into 96-well plates coated with 0.3% gelatin using a FACSAria-II cell sorter (BD Biosciences). Fast-growing wells (containing >1000 cells/well) were trypsinized and expanded to 24-well cell culture plates after 7-10 days. Culture medium was replaced daily. Genomic DNA from clones was isolated using DirectPCR (Tail) reagent (Viagen, catalog no. 102-T) and *Vcan* exon 2 and exon 3 were amplified using Phusion *Taq* (NEB, catalog no. F530L) (see SI). Amplicons were excised from 2% agarose gels, purified using the QIAquick Gel Extraction kit (QIAGEN, catalog no. 28704) and cloned into pCR-Blunt II-TOPO vector using the Zero blunt PCR cloning kit (Life Technologies, Invitrogen, catalog no. K2800-40). Plasmid DNA was harvested from bacterial colonies for Sanger-sequencing to determine the precise mutations. Two independent *Vcan*-null mESC clones, D8 and F9, were established. Western blotting and immunostaining of 10-day old embryoid bodies from these lines (see below) confirmed *Vcan* inactivation. Embryoid body formation and induction of vascular sprouts is described in the **Online Supplement**.

### Immunostaining and fluorescence microscopy

Immunostaining of E 9.5 and E 8.5 yolk sac was carried out on 30 μm thick vibratome sections [22] or paraffin-embedded 7 μm sections. Immunostaining of collagen-embedded embryoid bodies were carried out in 4-chamber cell culture slides (Fisher Scientific, catalog no. 354114). Confocal microscopy images of whole-mount mouse embryos and sections were acquired using a Leica TCS SP5 II multiphoton confocal microscope equipped with a 25X water immersion objective (Leica Microsystems, Wetzlar, Germany). For 3D-projection of whole-mount Z-stacks, the Volocity 3D imaging software was used (version 6.3, PerkinElmer, Inc., Waltham, MA) in maximum intensity projection method.

### Methylcellulose colony formation assay

Single cell suspensions of E8.5 embryos, yolk sacs or day-10 embryoid bodies were generated by incubation with trypsin for 10 minutes followed by disaggregation by pipetting with a 200 μL pipette tip until complete. 50,000 cells from each experimental group were transferred to a single 35 mm culture dish containing 1 mL of MethoCult GF M3534 culture medium (Methylcellulose medium with recombinant cytokines for mouse cells, Stem Cell Technologies, Vancouver, CA, catalog no. 03534) using a 3 mL syringe and 16-gauge needle, following the manufacturer’s protocol. Triplicate cultures from each genotype were incubated for 14 days in a humidified, 5% CO_2_, 37°C cell culture incubator. Blood colonies were counted using an inverted microscope. Aggregates with >50 cells were considered a colony-forming unit (CFU).

Additional details of reagents and procedures including primary antibodies used, RNAscope in situ hybridization, western blotting, transmission electron microscopy, and fluorophore-assisted carbohydrate electrophoresis (FACE) are described in the **Online Supplement**.

## Acknowledgments

This work was supported by the NIH-NHLBI Program of Excellence in Glycosciences award HL107147 (to S.S.A., R.J.M.,), NIH RF1 AG057579 (to Y.Y.) and by the Allen Distinguished Investigator Program, through support made by The Paul G. Allen Frontiers Group and the American Heart Association (to S.S.A), the David and Lindsay Morgenthaler Postdoctoral Fellowship and the Mark Lauer pediatric research grant (to S.N.). Purchase of the Leica SP8 confocal microscope was supported by NIH SIG grant 1S10RR026820-01. We thank Dr. David LePage and Dr. Ron Conlon at the CWRU transgenic core for R1 mES cells, Eric Schultz and Joseph Gerow of the LRI Flow Cytometry Core for mES cell sorting, Valbona Cali for FACE assays, Dr. Judy Drazba and Mei Yin of the LRI Imaging Core for guidance with confocal and electron microscopy and the Apte laboratory members for valuable discussions.

## Author contributions

S.N. and A.O conducted experiments. S.N., R.J.M., and S.S.A. conceived experiments and interpreted the data. S.N. and S.S.A. wrote the paper. All authors read and edited the paper.

## SUPPLEMNTAL FIGURES AND TABLE

**Figure 1, Supplement 1:**
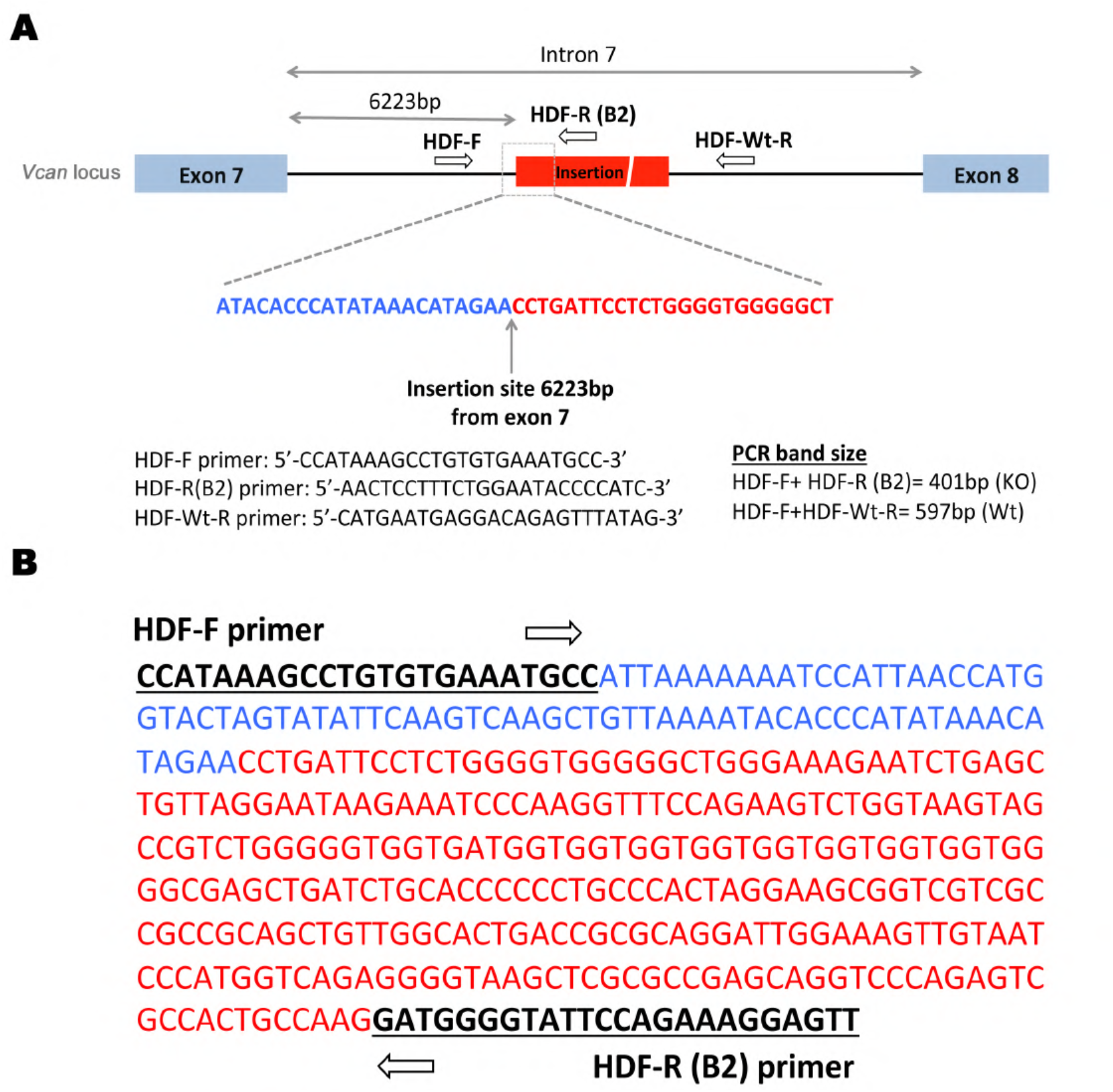
Characterization of the *Vcan^hdf^* mutation and genotyping strategy. **(A)** *Vcan* locus, showing the retroviral insertion site in intron 7. *Vcan* exons are indicated in blue and the insertion in red. Primer sequences and amplicons used for genotyping are shown at the bottom. **(B)** Sequence of the 401bp *Vcan^hdf^* amplicon and the primer sequences used (underlined). The *Vcan* intron 7 sequence is in blue text and sequence corresponding to the retroviral insertion is in red text.

**Figure 1, Supplement 2:**
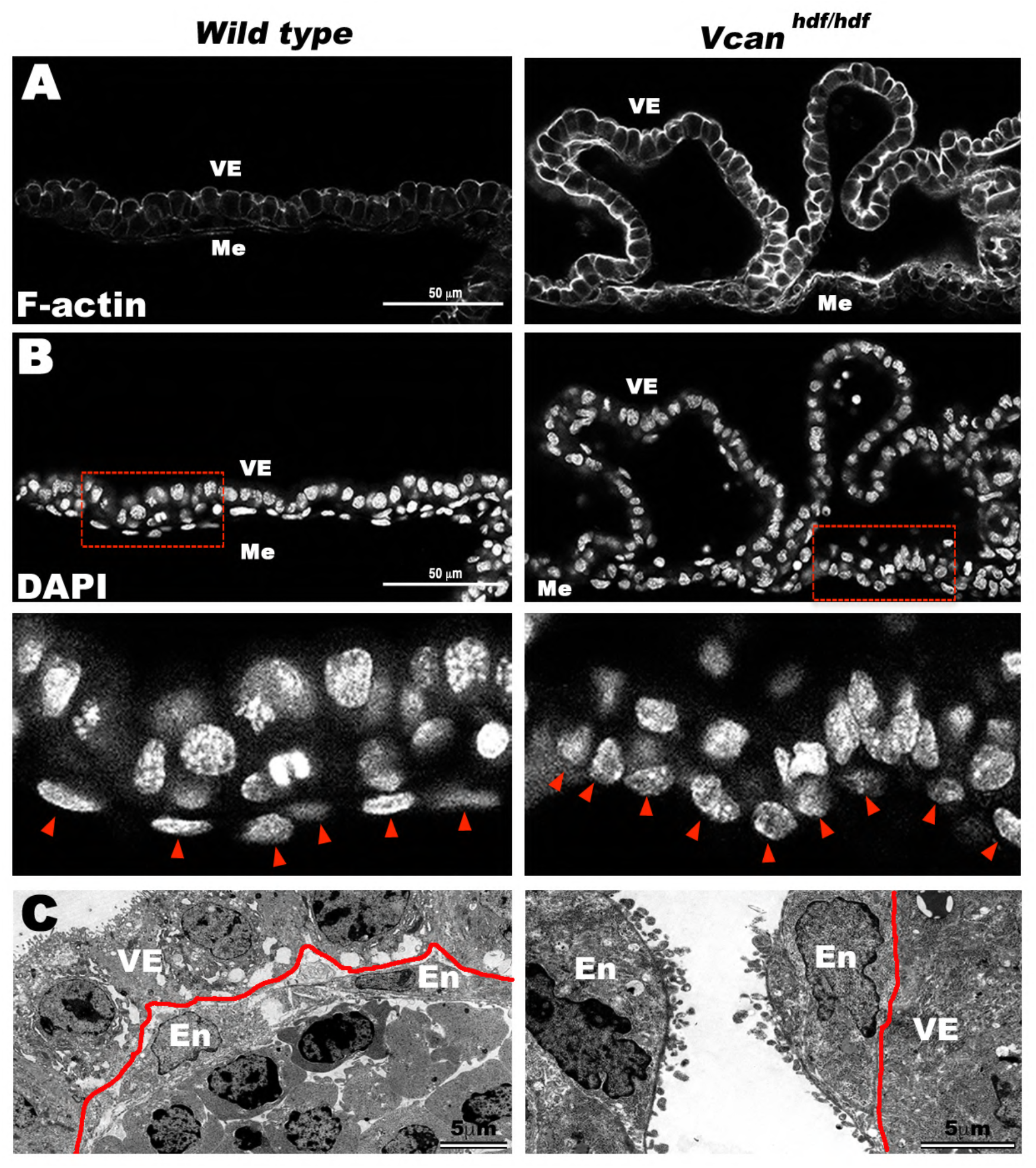
Severe disorganization and cellular changes in *Vcanhdf/hdf* yolk sacs. **(A)** E9.5 *Vcanhdf/hdf* yolk sacs show dramatically increased F-actin staining. **(B)** DAPI staining shows altered nuclear morphology and disrupted cellular organization of the *Vcanhdf/hdf* yolk sac mesoderm (Me) and mesothelium and (with **A**) separation from visceral endoderm (VE). High magnification of the boxed areas (red line) in the upper panels of **B** are shown below. Arrowheads identify the nuclei of yolk sac mesothelium, which are rounded in *Vcanhdf/hdf* yolk sac. **(C)** Transmission electron microscopy (TEM) images of wild type and *Vcanhdf/hdf* yolk sacs showing endothelium-lined blood islands in the wild type and detached vascular endothelial cells in the *Vcanhdf/hdf* yolk sac. The red line marks the basement membrane of the visceral endoderm, En=vascular endothelium. Scale bars in **A-B**= 50μm, 5μm in **C**.

**Figure 4, Supplement 1:**
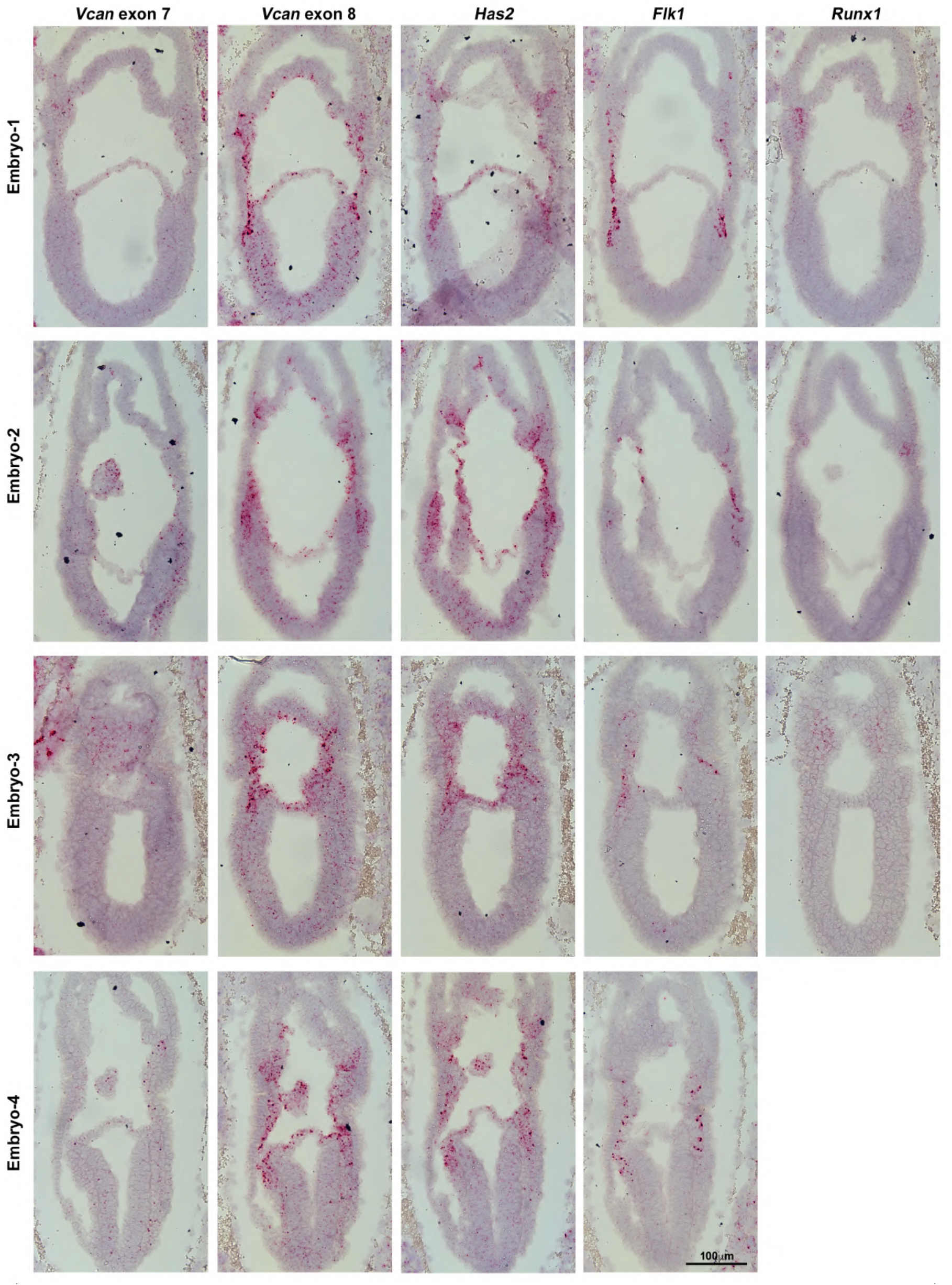
RNAScope *in situ* hybridization of *Vcan* exon 7, *Vcan* exon 8, *Has2*, *Flk1* and *Runx1* probes to consecutive sections from four individual E7.5 wild type embryos. Hybridization signal is red, sections were counterstained with hematoxylin (blue). Scale bar = 100μm.

**Figure 4, Supplement 2:**
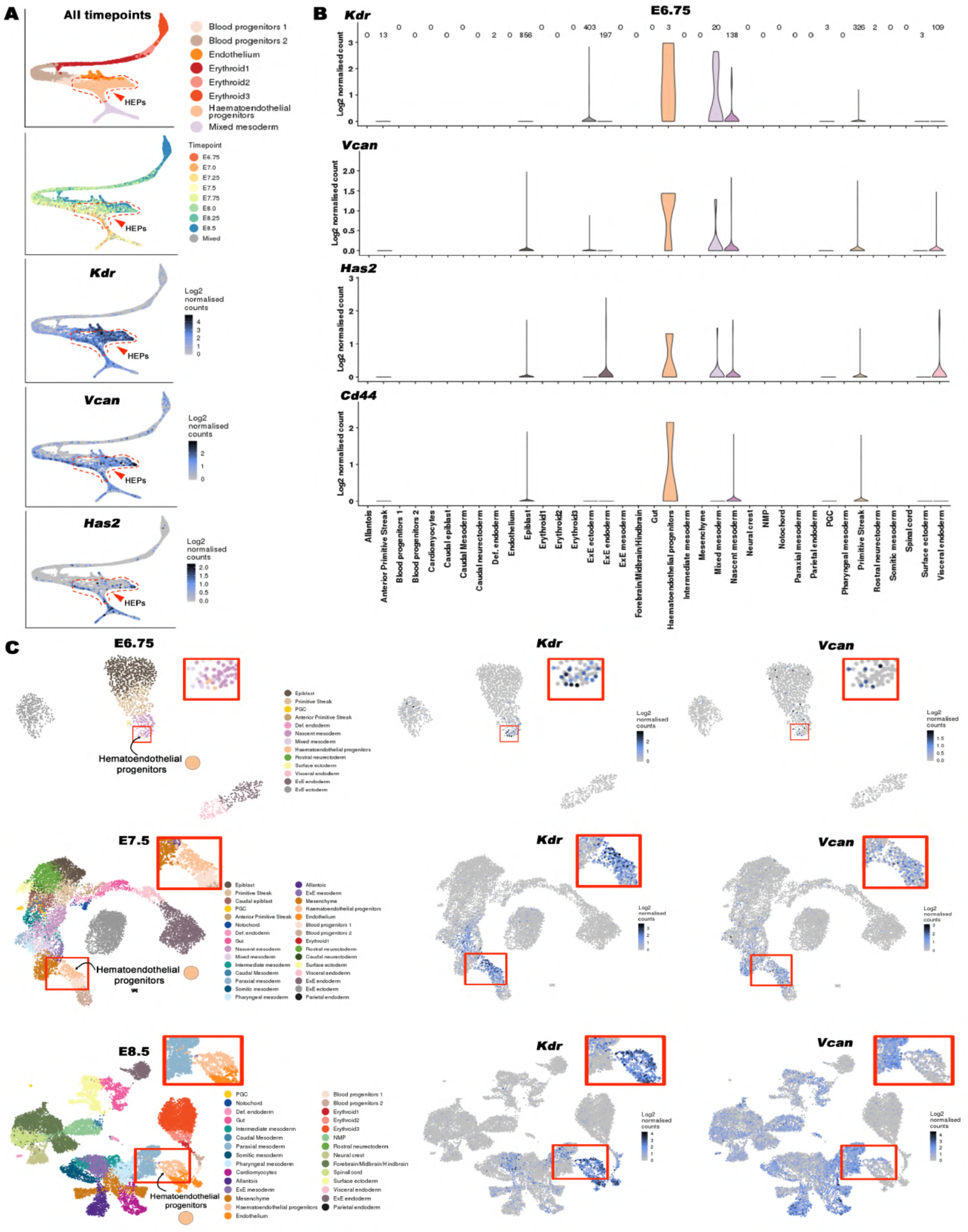
Data extracted from a published single cell transcriptome atlas of mouse gastrulation illustrates specific *Vcan* and *Has2* expression by *Kdr*-expressing hematoendothelial cells. **(A)** Force-directed graph layout of blood-related lineages showing the lineage-specific and time-resolved expression of *Kdr, Vcan* and *Has2*. The stippled red line encircles hematoendothelial progenitors. **(B)** Violin plots of *Kdr, Vcan*, *Has2* and *Cd44* expression by cells of various lineages from E6.75 embryos. Note strong co-expression of these genes by single hematoendothelial progenitors. **(C)** Uniform manifold approximation and projection (UMAP) plots showing overlap between single cell *Kdr* and *Vcan* expression at different gestational ages. High magnification inserts of hematoendothelial progenitors are indicated within red boxes.

**Figure 4, Supplement 3:**
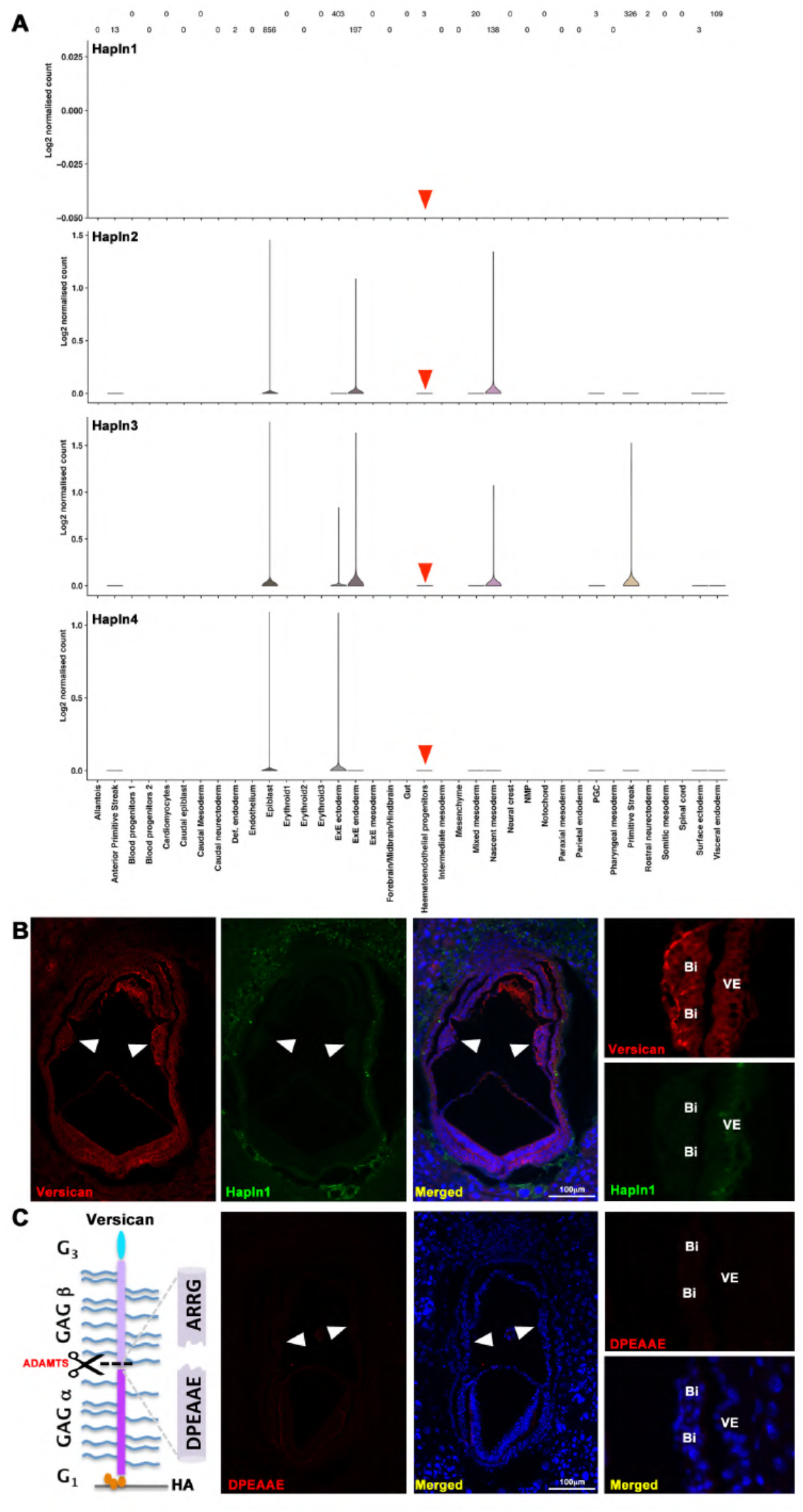
Data extracted from a published single cell transcriptome atlas of mouse gastrulation illustrates lack of link protein gene expression by hematoendothelial progenitors and versican cleavage in blood islands. **(A)** Violin plots for *Hapln1-4* expression by cells of various lineages from E6.75 embryos. No expression of *Hapln1-4* is detected in hematoendothelial progenitors (red arrowhead) at this stage. **(B)** HAPLN1 staining was not detected in E7.5 blood islands marked by versican staining (white arrowheads). **(C)** ADAMTS-cleaved versican staining (anti-DPEAAE) is undetectable in E7.5 blood islands (white arrowheads). Scale bars = 100μm in **B** and **C**.

**Figure 5, Supplement 1:**
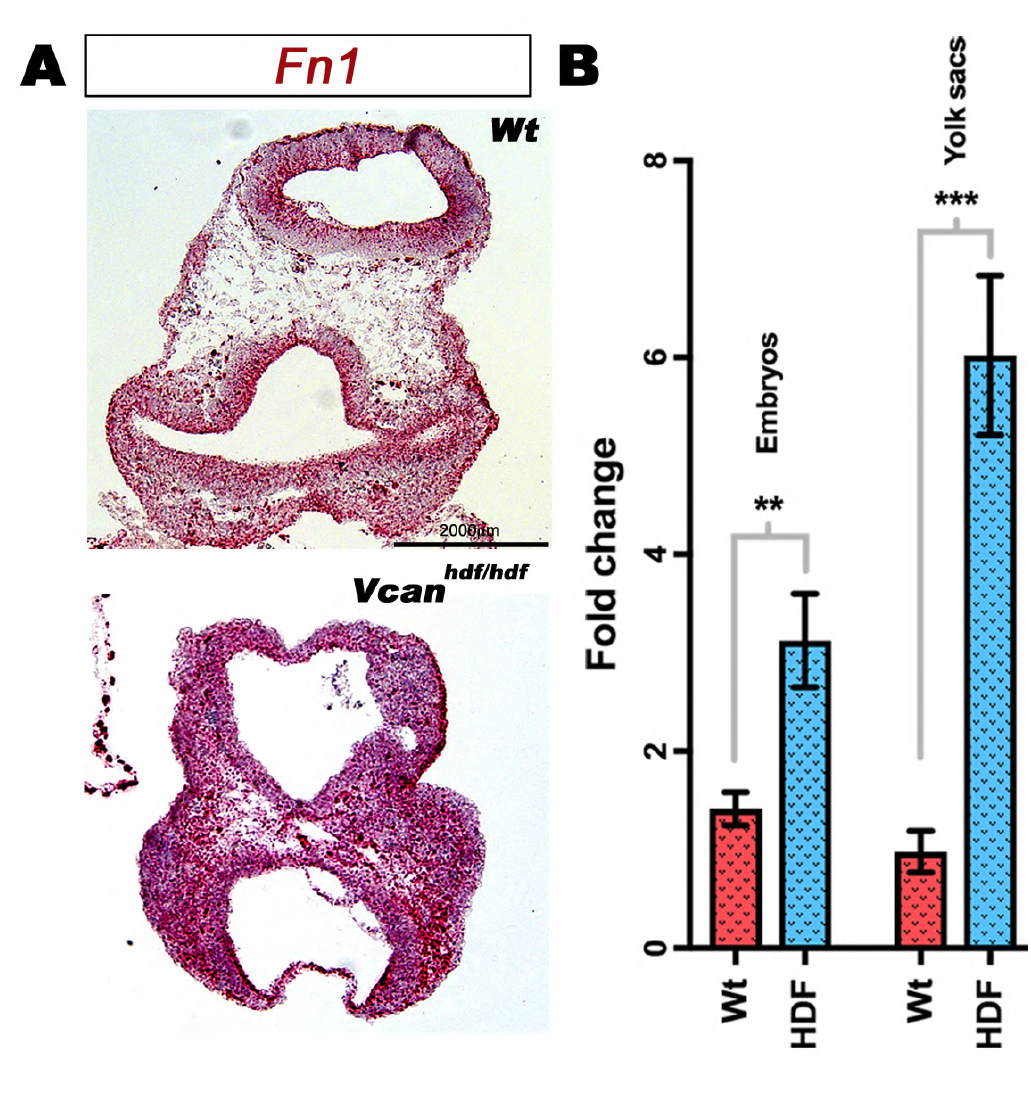
Increased *Fn1* transcription in *Vcan^hdf/hdf^* embryos. **(A)** RNAscope *in-situ* hybridization of E9.5 wild type and *Vcan^hdf/hdf^* embryos showing stronger *Fn1* RNA expression in the mutant embryo. **(B)** qRT-PCR analysis of *Fn1* transcript in wild type and *Vcan^hdf/hdf^* embryos and yolk sacs showing significantly increased expression in the mutant. (n=3 embryos and yolk sacs each genotype, error bars= S.E.M., **, p<0.01; ***, p<0.0001). Scale bars = 100μm in **A**.

**Figure 6, Supplement 1:**
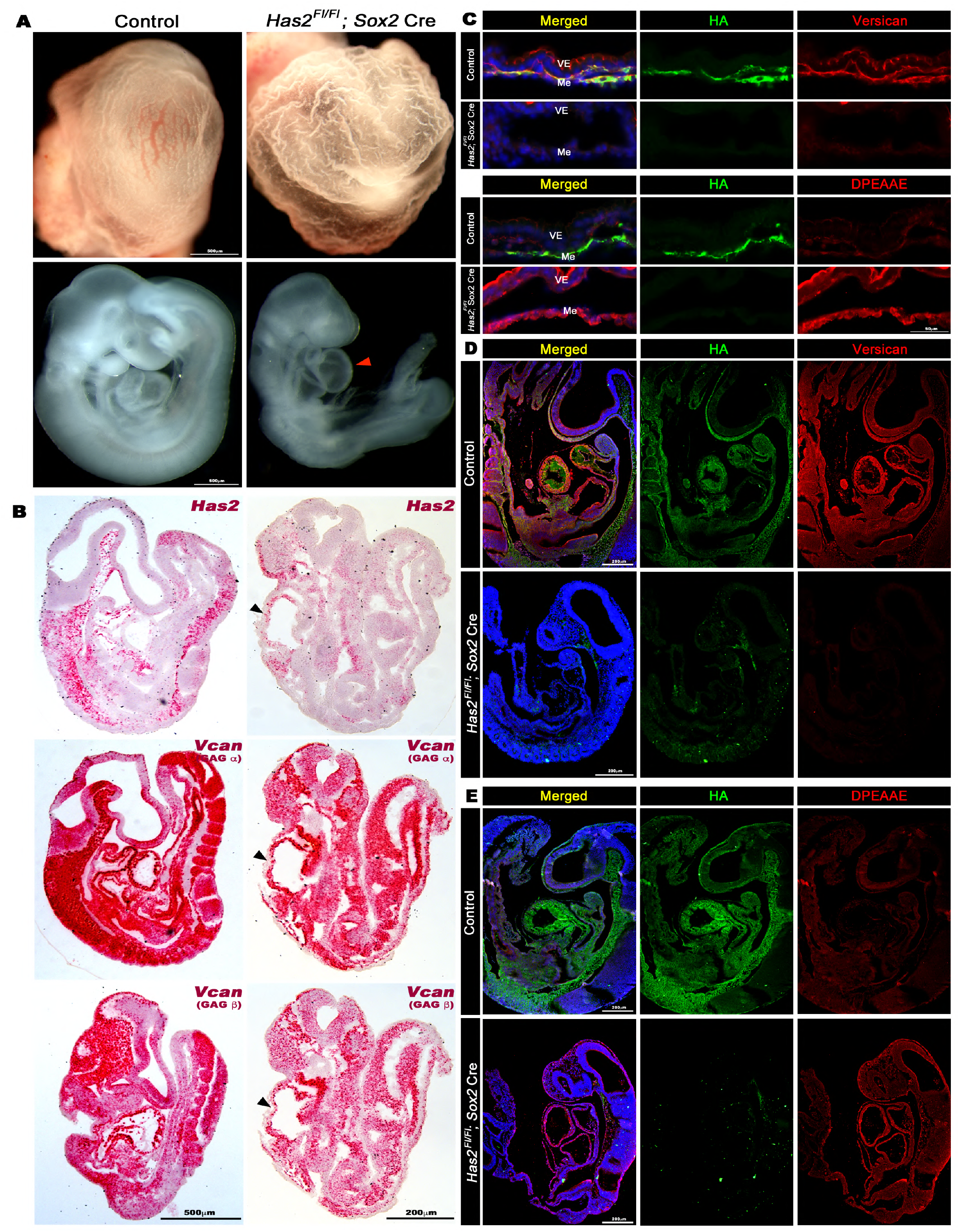
Loss of yolk sac vasculature, HA and versican staining in *Has2* conditionally deleted embryos. **(A)** Whole mount images of *Has2^Fl/Fl^; Sox2*Cre yolk sacs and embryos showing loss of yolk sac vasculature and heart defect (red arrowhead) compared to control littermates (N=8 each genotype). **(B)** RNAScope in situ hybridization for *Has2, Vcan* exon 7 (GAG *α*) and exon 8 (GAG *β*) shows residual *Has2* transcript and slightly reduced *Vcan* transcript labeling in *Has2^Fl/Fl^; Sox2* Cre embryos (n=4 embryos each probe). **(C)** *Has2^Fl/Fl^; Sox2* Cre yolk sacs show loss of HA and versican staining (upper panels) and increased versican catabolism is revealed by DPEAAE staining (lower panels) (N=4 each group). **(D)** *Has2^Fl/Fl^; Sox2* Cre embryos show loss of HA and versican staining (N=3 each group). **(E)** *Has2^Fl/Fl^; Sox2* Cre embryos show increased DPEAAE staining (N=3 each group).

**Figure 7, Supplement 1:**
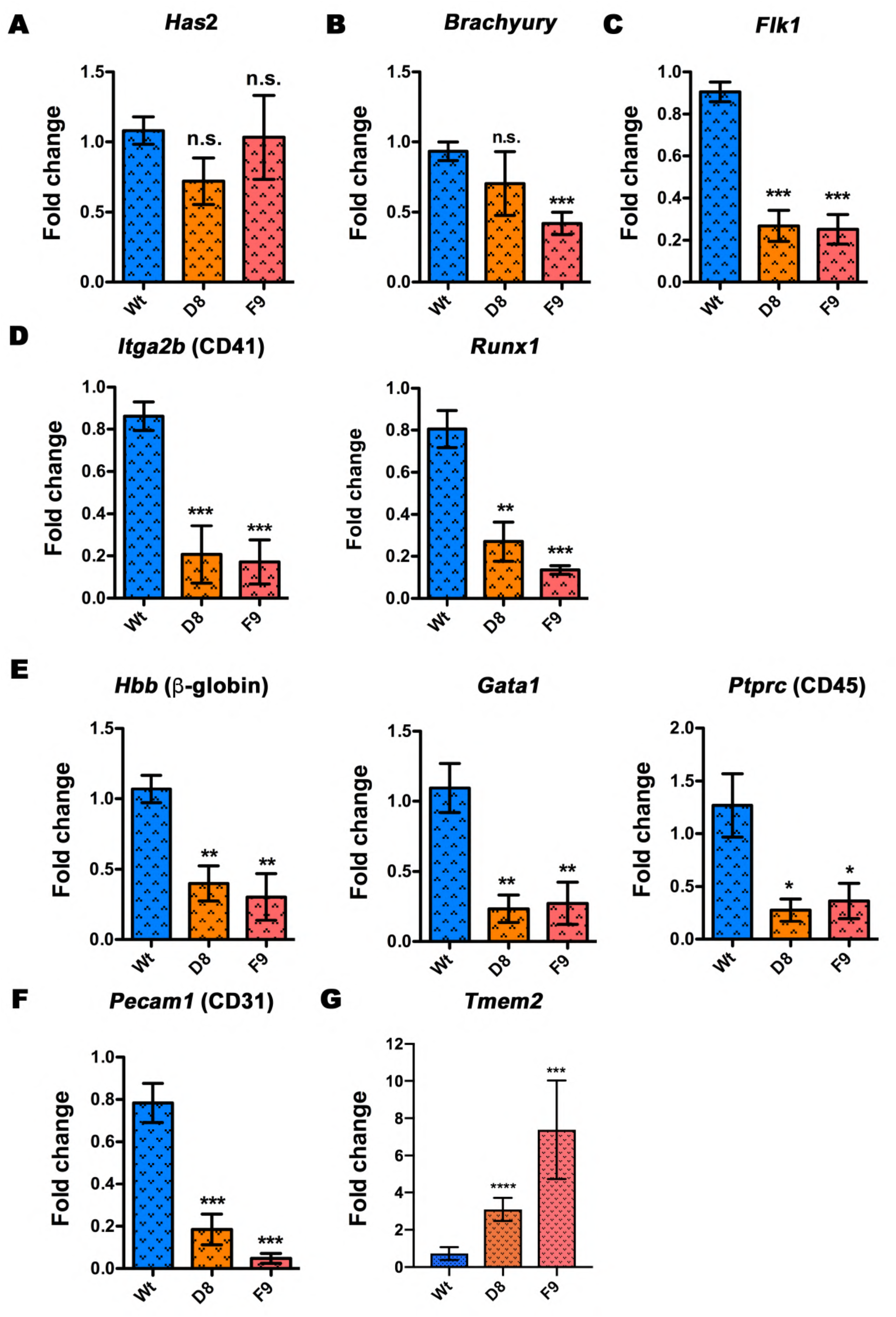
Formation of blood and vascular lineages is impaired in *Vcan-*null embryoid bodies. **(A-F)** 10-day differentiated embryoid bodies from three independent experiments were analyzed by qRT-PCR. *Has2* transcription in embryoid bodies was unaffected by the loss of versican. mRNA for the mesoderm marker *Brachyury* was significantly lower in F9 but not in D8 embryoid bodies contrasting with *Vcan^hdf/hdf^* embryos (main Fig. 3E). Expression of the hematovascular progenitor marker *Flk1,* blood lineage commitment progenitor markers *Itga2b* (CD41) and *Runx1*, differentiated blood markers *Hbb* (*β*-globin), *Gata1* and *Ptprc* (CD45) and endothelial marker *Pecam1* (CD31) was reduced in *Vcan*-null embryoid bodies (N=3 independent batches of 10 day embryoid bodies from each genotype, error bars= S.E.M.,*, p<0.05; **, p<0.01; ***, p<0.0001).

**Figure 8, Supplement 1:**
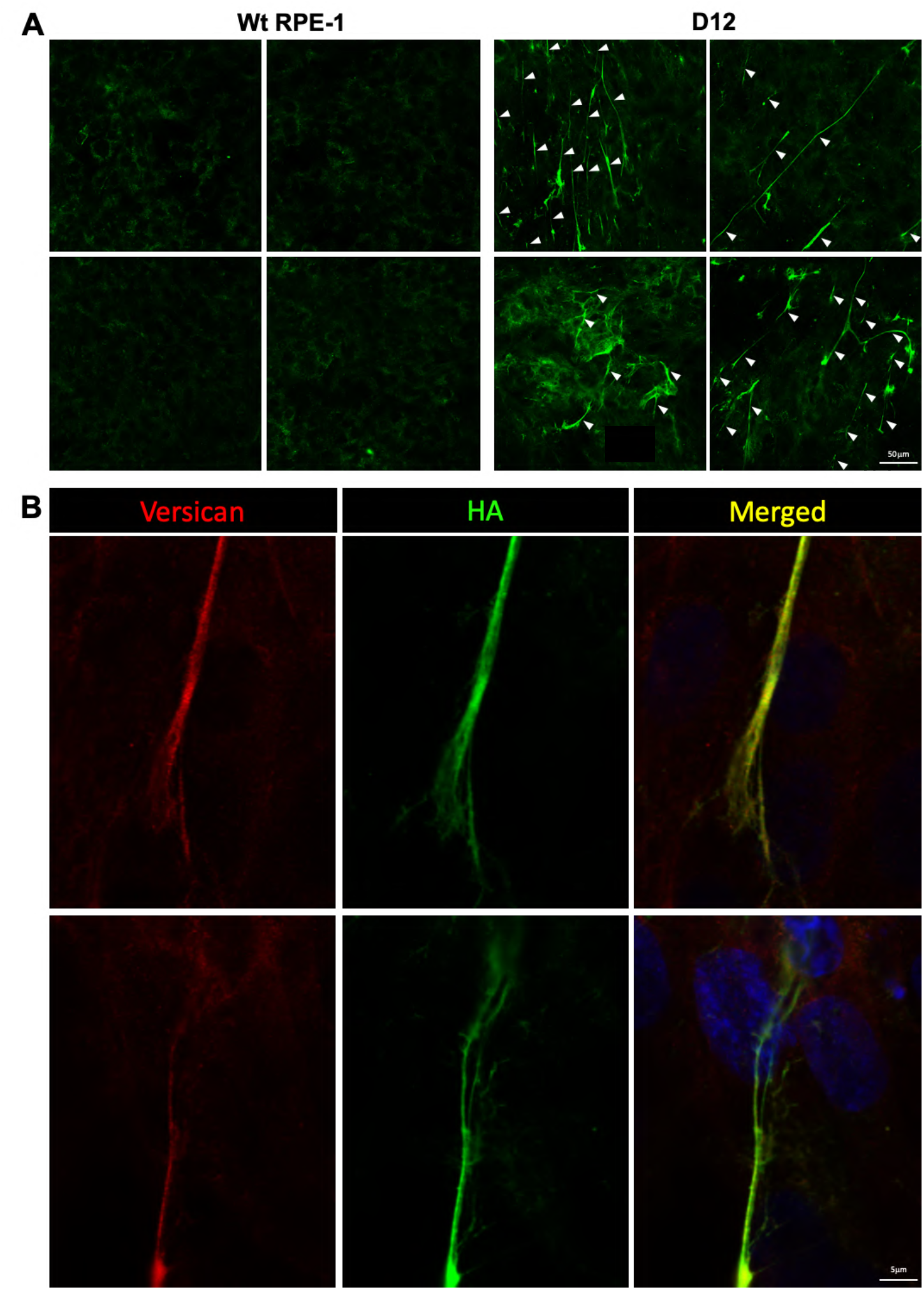
Formation of hyaluronan and HA cables in *ADAMTS9* deficient RPE-1 cells in 25mM glucose. **(A)** Four independent representative areas of wild type RPE-1 and *ADAMTS9* deficient RPE-1 (D12) cell cultures stained with HAbp showing HA accumulation and numerous long HA-cables present in D12 cultures, but not parental RPE1 cells (white arrowheads). **(B)** High-magnification confocal images showing two examples of long HA cables (green) decorated with versican (red). Scale bars = 50μm in **A** and 5μm in **B**.

**Figure 8, Supplement 2:**
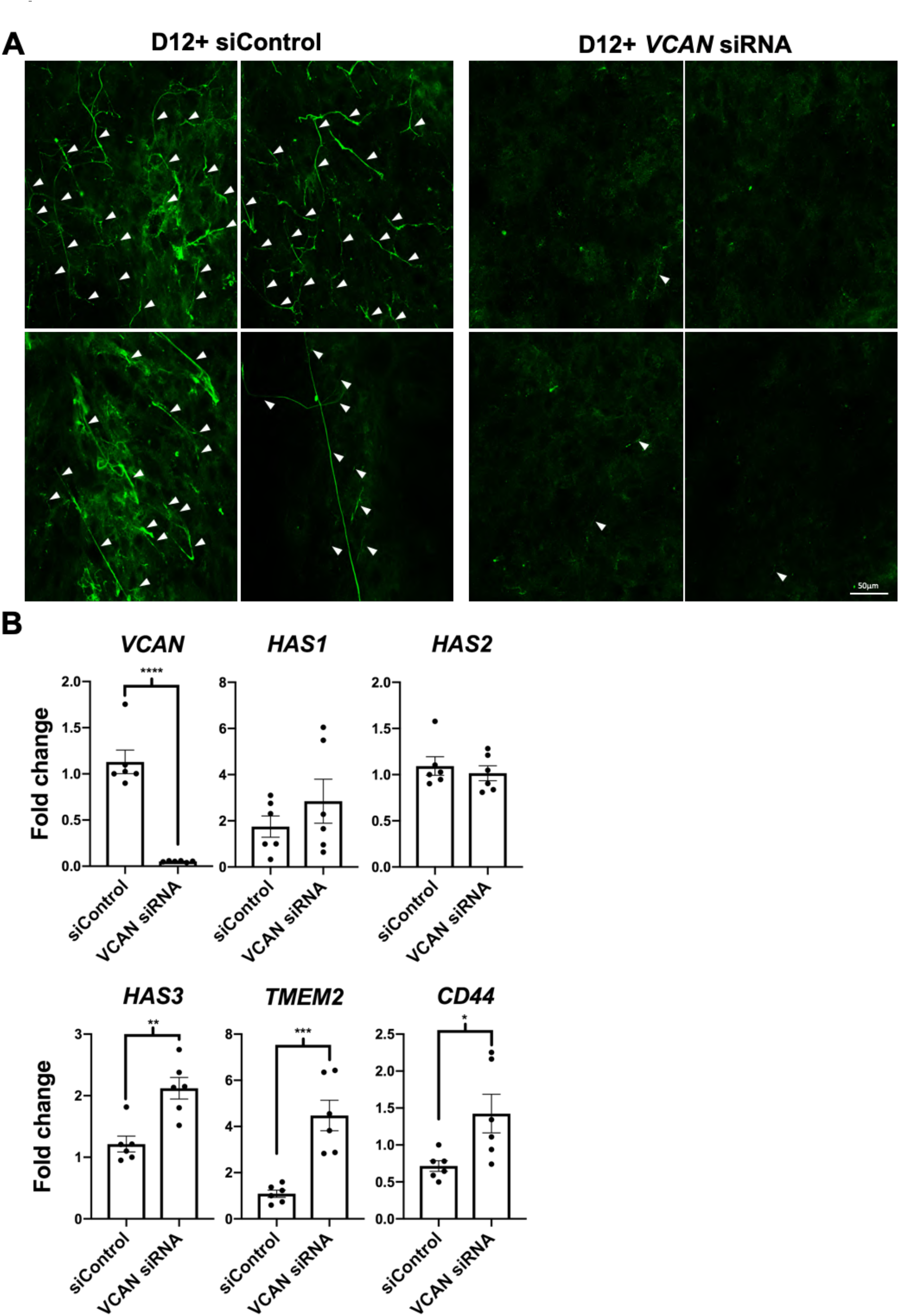
Loss of HA and HA cables in *ADAMTS9*-deficient RPE-1 cells upon *VCAN* siRNA treatment. **(A)** Four independent representative areas of *ADAMTS9*-deficient RPE-1 (clone D12) cell cultures transfected with control siRNA (siControl) or *VCAN* siRNA, stained with HAbp showing loss of HA after *VCAN* knockdown, whereas HA-cables persist in siControl D12 cells (white arrowheads). **(B)** qRT-PCR of siControl and *VCAN* siRNA treated D12 cells showing decreased *VCAN* transcript levels upon siRNA treatment and upregulation of *TMEM2* expression (n=3 independent siRNA treatment experiments, error bars= S.E.M., *, p<0.05; **, p<0.01; ***, p<0.001; ****, p<0.0001). Scale bar = 50μm in **A**.

**Figure 8, Supplement 3:**
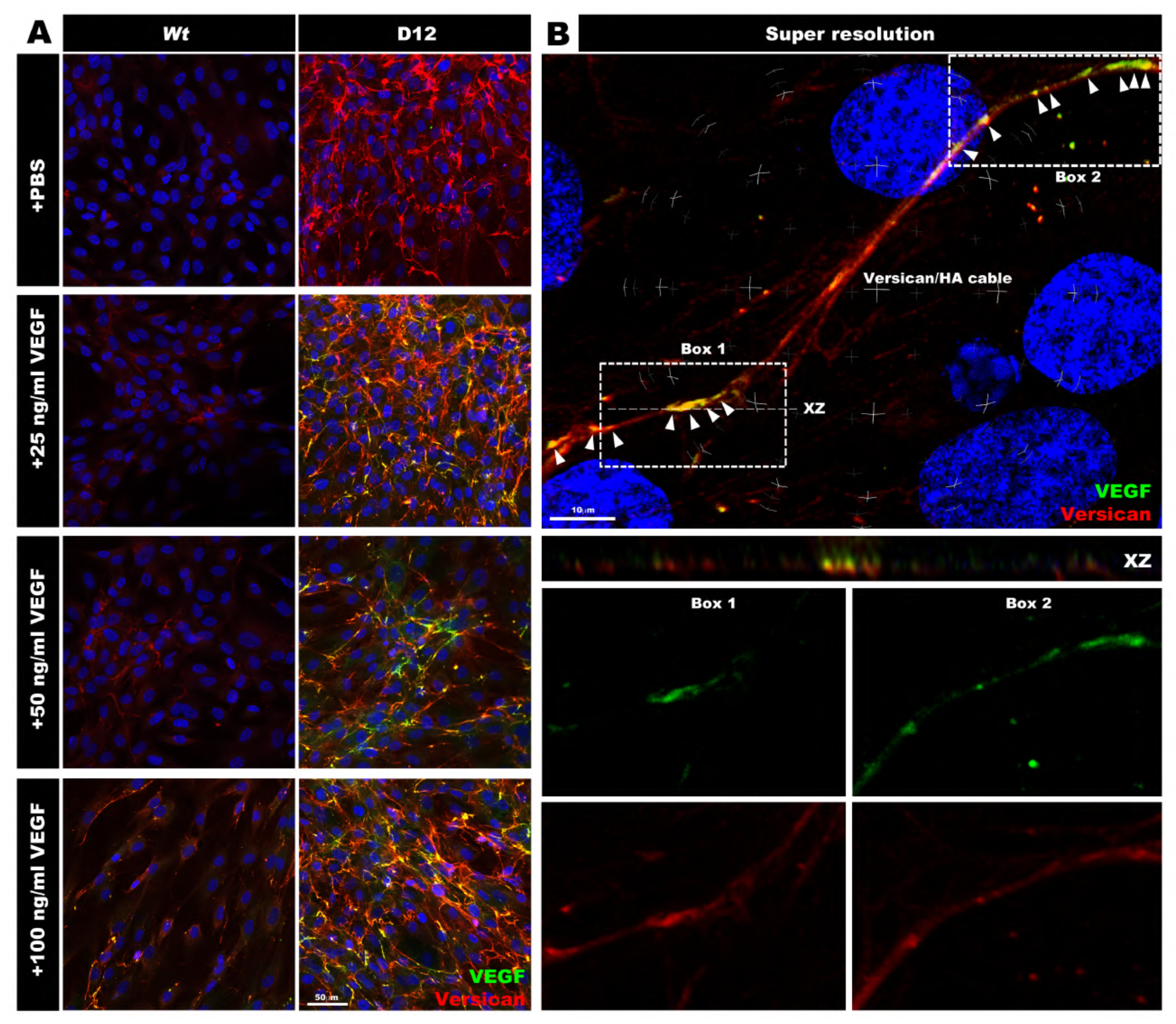
VEGF_165_ co-localizes with versican in extracellular matrix of RPE1 D12 cells in a dose-dependent manner. **(A)** Wild type RPE-1 and *ADAMTS9*-deficient RPE-1 (D12) cells treated with PBS or increasing concentrations of recombinant biotinylated VEGF_165_ (green) showing dose-dependent VEGF staining intensity and colocalization with versican (red) in D12 cells. **(B)** Super-resolution confocal images of a long versican/HA cable showing patches of VEGF_165_ (green) deposited along the versican stained cable (red). Nuclei are stained blue with DAPI. The upper merged panel shows some strongly co-staining regions (arrowheads), whereas the center panel shows co-staining in the X-Z plane. The lower panels show the single-color images of the areas marked as Box 1 and Box 2 in the upper panel. Scale bars = 50μm in **A** and 10μm in **B**.

**Figure 8, Supplement 4:**
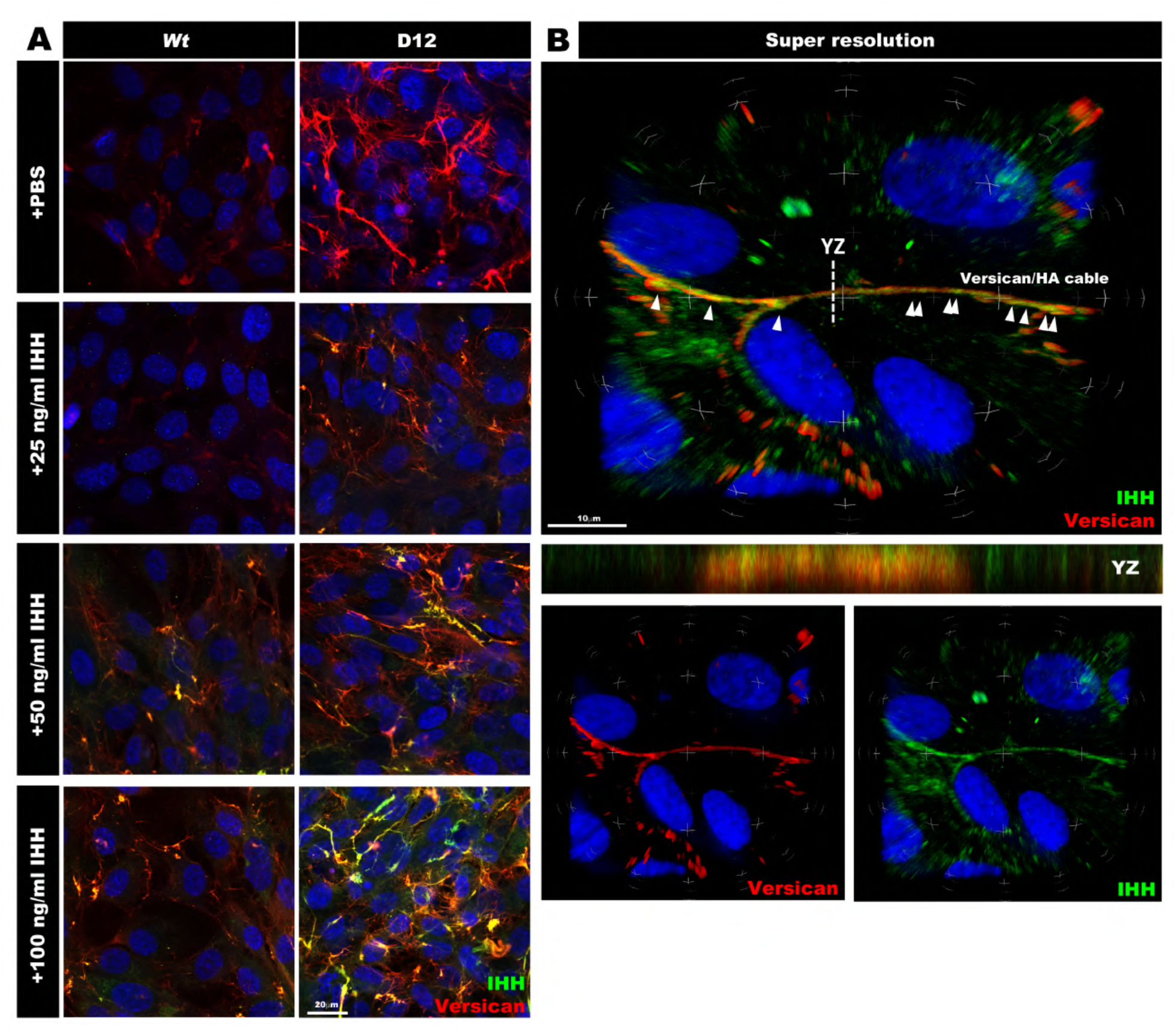
Ihh co-localizes with versican in extracellular matrix in a dose-dependent manner. **(A)** Wild type RPE-1 and *ADAMTS9* deficient (D12) cells treated with PBS or increasing concentrations of recombinant Ihh N-terminal domain (green) showing dose-dependent deposition and increased co-staining (yellow) with versican (red) in D12 cells. Nuclei are stained blue with DAPI. **(B)** Super-resolution confocal image of a versican-HA stained cable showing Ihh staining (green) along a versican stained cable (red). Strongly co-stained regions are indicated by arrowheads. The center panel shows the Y-Z plane. Bottom panels show single-color images of the area imaged in the upper panel. Scale bars = 20μm in **A** and 10μm in **B**.

**Supplemental Table 1.**
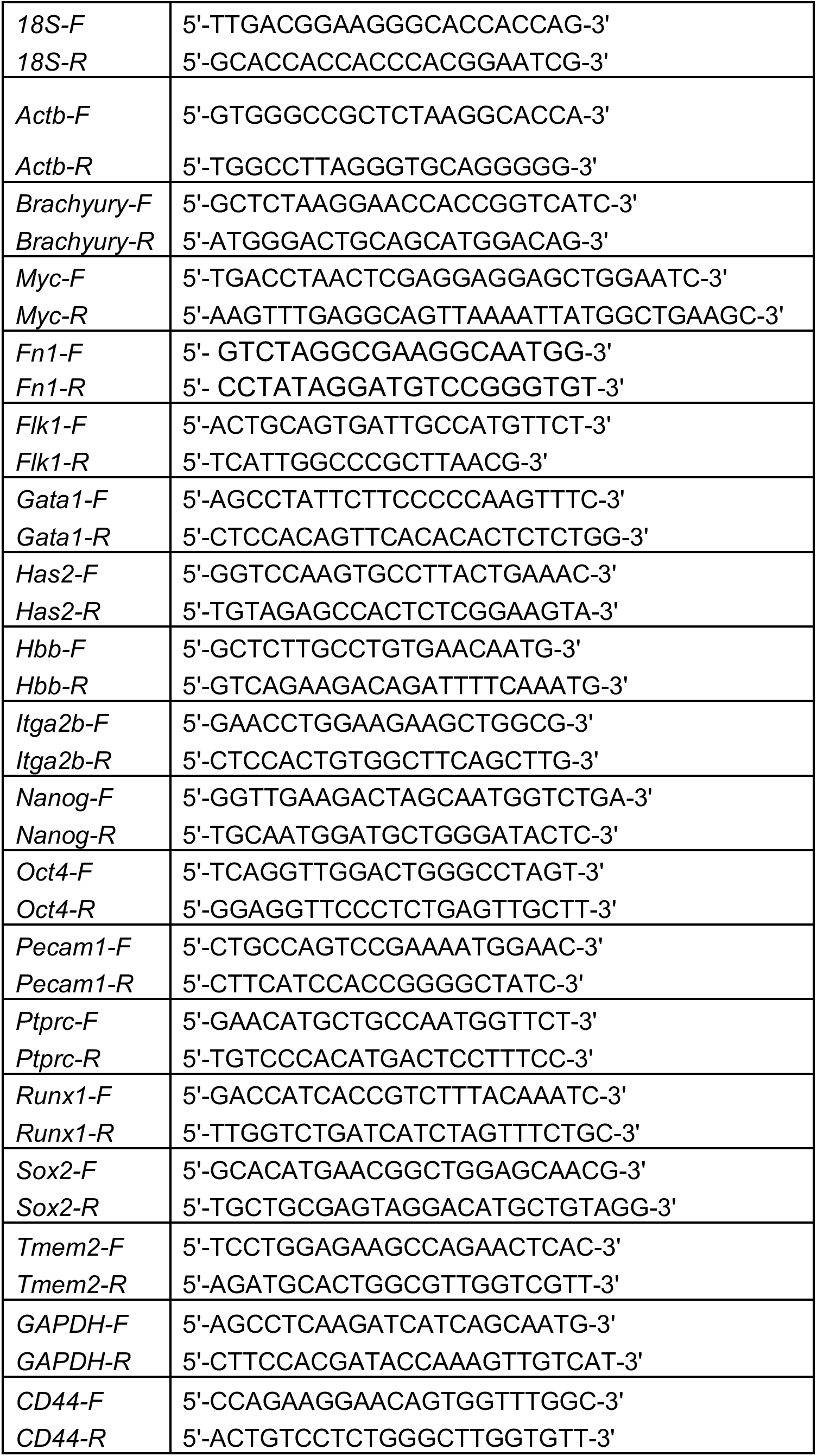
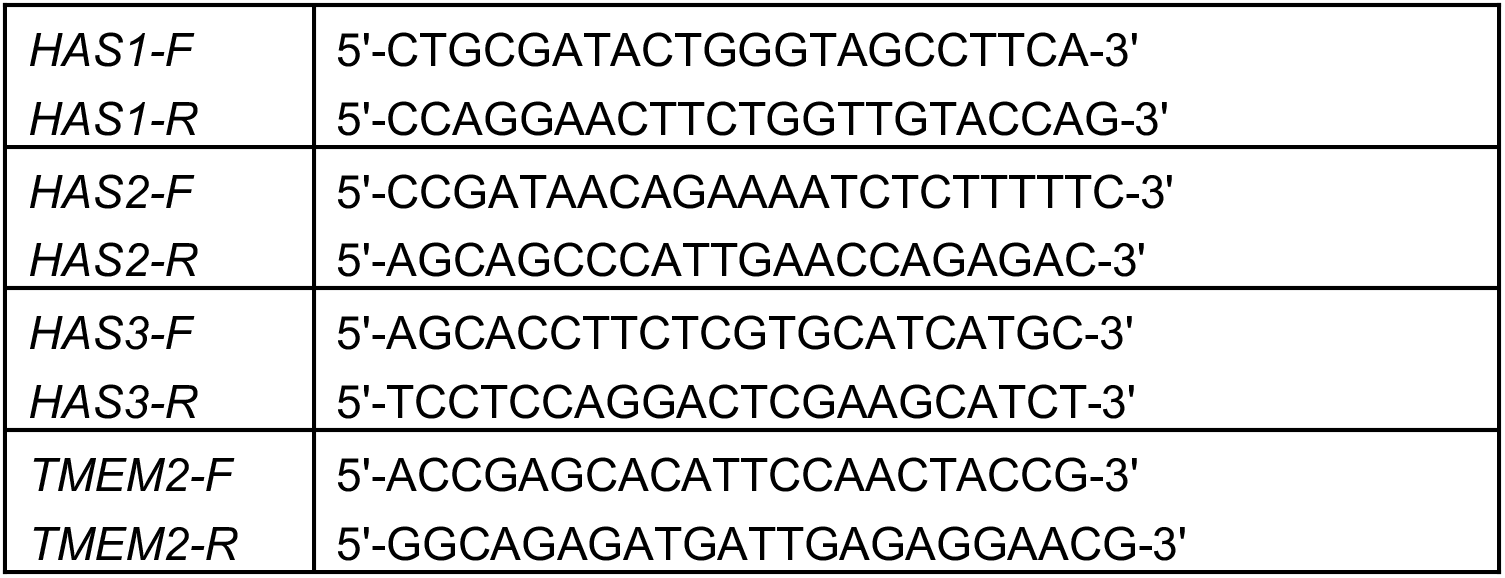
Oligonucleotide primers used for qRT-PCR.

## SUPPLEMENTAL METHODS

### Mouse genotyping

*Vcan*^Tg(Hoxa1)1Chm^ (*Vcan*^hdf^) genotyping is described in Figure-1, Supplement-1. *Has2^flox^* genotyping was carried out using the following primers; forward primer: 5**^’^**-TGCAGAATTTAGGGGCGAATTGGGAGCTAA-3**^’^**, reverse primer: 5**^’^**-ATGAGGTTAGAGATTAGCAAGACTGAGTTC-3**^’^** which results in a 441bp band for wild type and a 550bp band for the floxed allele. Sox2-Cre mice were genotyped using the following three-primer combination, primer-1: 5**^’^**-CTTGTGTAGAGTGATGGCTTGA-3**^’^**, primer-2: 5**^’^**-TAGTGCCCCATTTTTGAAGG-3**^’^**, primer-3: 5**^’^**-CCAGTGCAGTGAAGCAAATC-3**^’^** which results in a 207bp band for wild type and a 165bp band for Sox2-Cre. The *Has2* null allele was genotyped using forward primer: 5**^’^**-CTTGAACCTTGAGTGTGCCATTTTGTAGTC-3**^’^**, reverse primer: 5**^’^**-CATTCTTGTTTTGAAGTTTGTTTCCTTGAC-3**^’^** which results in a 346bp band.

### Immunostaining and fluorescence microscopy

Immunostaining of E9.5 and E8.5 yolk sac was carried out on 30μm thick vibratome sections [1] or paraffin-embedded 7 μm sections. Immunostaining of cultured RPE-1 cells were carried out in 8-chamber cell culture slides (Fisher Scientific, catalog no. 354118). Immunostaining of collagen-embedded embryoid bodies were carried out in 4-chamber cell culture slides (Fisher Scientific, catalog no. 354114). Confocal microscopy images of whole mount mouse embryos and sections were acquired using a Leica TCS SP5 II multiphoton confocal microscope equipped with a 25X water immersion objective (Leica Microsystems, Wetzlar, Germany). For 3D-projection of whole mount Z-stacks, the Volocity 3D imaging software was used (version 6.3, PerkinElmer, Inc., Waltham, MA) in maximum intensity projection method. Confocal and super resolution microscopy of RPE-1 cells were carried out using a Leica TCS SP8 confocal microscopy equipped with Huygens deconvolution (HyVolution) capability as previously described [2].

### Methylcellulose colony formation assay

Single cell suspensions of E8.5 embryos, yolk sacs or day-10 embryoid bodies (EBs) were generated by incubation with trypsin for 10 minutes followed by disaggregation by pipetting with a 200 μL pipette tip until complete. 50,000 cells from each experimental group were transferred to a single 35 mm culture dish containing 1 mL of MethoCult GF M3534 culture medium (Methylcellulose medium with recombinant cytokines for mouse cells, Stem Cell Technologies, Vancouver, CA, catalog no. 03534) using a 3 mL syringe and 16-gauge needle, following the manufacturer’s protocol. Triplicate cultures from each genotype were incubated for 14 days in a humidified, 5% CO_2_, 37°C cell culture incubator. Blood colonies were counted using an inverted microscope. Aggregates with >50 cells were considered a colony-forming unit (CFU).

### Primary antibodies and dilutions

For staining whole mount embryos, yolk sacs, vibratome sections and cell culture chamber slides the following antibodies, staining reagents and dilutions were used: Rabbit polyclonal anti-mouse versican GAG*β* domain (Millipore-Sigma, catalog no. AB1033) 1:200; Rabbit polyclonal anti-versican cleavage site (anti-Vc) [3] 1:400; Rabbit polyclonal anti-versican V_0_/V_1_ neo cleavage antibody (DPEAAE) (Thermo Fisher, catalog no. PA1-1748A) 1:400; biotinylated hyaluronan binding protein (HAbp) (Millipore-Sigma, Calbiochem, catalog no. 385911) 1:100; rabbit polyclonal anti-fibronectin (Abcam, catalog no. Ab2413) 1:200; rabbit polyclonal anti-collagen-IV (Rockland antibodies and assays, catalog no. 600-401-106) 1:400; rat monoclonal anti-mouse Flk1 (clone Avas12, Thermo Fisher, catalog no. 17-5821-81) 1:200; hamster anti-mouse CD31 (Milipore-Sigma, catalog no. MAB1398Z) 1:400; FITC-conjugated anti-mouse CD41 (Biolegend, catalog no.133904) 1:100; mouse monoclonal Cy3 conjugated smooth muscle *α*-actin (Sigma-Aldrich, catalog no. C6198) 1:600;Alexa Fluor-568 phalloidin (Life Technologies, catalog no. A12380) 1:500; Streptavidin-FITC (Invitrogen, catalog no. SA1001) 1:400; Goat anti-mouse Ihh N-terminus (R&D Systems, catalog no. AF1705) 1:400; Mouse anti-Hapln1 (Link protein) (DSHB, 9/30/8-A-4-C) 1:100. All primary antibodies were diluted in 5% normal goat serum in PBST (PBS+ 0.1% Tween 20) and incubated overnight at 4°C. Alexa488, 568 or 647-conjugated secondary antibodies against the corresponding species, or streptavidin conjugated Alexa488 or 568-labeled antibodies for HAbp detection were purchased from Invitrogen and were used at 1:400 dilution at room temperature for 2-3hrs. Slides were photographed using an Olympus BX51 upright microscope (Olympus, Center Valley, PA) connected to a Leica DFC7000T camera and Leica Application Suite v4.6 imaging software (both from Leica, Wetzlar, Germany).

### Quantitative real time RT-PCR (qRT-PCR) analysis

RNA from whole embryos, dissected yolk sacs, cultured cells or EBs was extracted using TRIzol reagent (ThermoFisher Scientific, catalog no. 15596026) according to manufacturer recommendations. 2 μg of RNA from each sample (embryo, yolk sac or EB pool) was used for cDNA synthesis using the high capacity cDNA synthesis kit (Applied Biosystems, Thermo Fisher Scientific, catalog no. 4368814). qRT-PCR was carried out using a CFX96 Touch Real-Time PCR detection system (Bio-Rad Laboratories) with the Bullseye EvaGreen qPCR mix (MIDSCI, catalog no. BEQPCR-S), using primer pairs listed in the Supplemental Table-1. 18s ribosomal RNA or *Actb* RNA were used for normalization of mouse genes while *GAPDH* was used for normalizing human gene expression and relative expression of genes was calculated by the *ΔΔ*Ct quantification method. An unpaired, two-tailed Student t-test was used to determine statistical significance.

### RNAscope *In-situ* hybridization and RNAseq database mining

4% PFA fixed mouse embryos and yolk sacs were paraffin embedded and 7 μm thick sections were collected immediately prior to in-situ hybridization. All RNAscope in situ probes and reagents were from Advanced Cell Diagnostics. The probes were: *Vcan* exon 7 (GAG-*α*) (catalog no. 428311), *Vcan* exon 8 (GAG-*β*) (catalog no. 428321), *Has2* (catalog no. 465171), *Flk1 (Kdr)* (catalog no. 414811) and *Runx1* (catalog no. 406671). In situ hybridization was carried out following the user manual for RNAScope 2.5 HD Red detection kit using the RNAscope HybEZ oven. Slides were photographed as described above.

A database containing recent analysis of 116,312 single cells (Pijuan-Sala B, et al., 2019) obtained from gastrulating mouse embryos spanning E6.5 to E8.5 was searched at specific gestational ages for expression of *Vcan, Kdr, Has2* and *Cd44*. Individual data panels were generated by the web portal for the deposited scRNAseq data accessed via: https://marionilab.cruk.cam.ac.uk/MouseGastrulation2018/

### Western blotting

7.5% reducing SDS-PAGE was used for western blotting. Embryoid bodies were lysed in PBS containing 1% Tween 20 supplemented with a protease inhibitor cocktail (Roche, catalog no. 11873580001). Supernatants obtained after centrifugation at 1200 rpm for 10 minutes were treated with BSA free *Proteus vulgaris* chondroitinase ABC (Sigma-Aldrich, catalog no. 3667) for 2 h at 37°C. Samples were boiled in Laemmli sample buffer (0.375% mM Tris.HCL, 9% SDS, 50% glycerol, 0.03% bromophenol blue, with 9% (v/v) *β*-mercaptoethanol) for 10 minutes followed by SDS-PAGE. Nitrocellulose membranes with transferred proteins were blocked in LI-COR Odyssey PBS blocking buffer (LI-COR Biosciences, Lincoln, NE catalog no. 927-40000) for 30 minutes in RT. Primary antibodies rabbit anti-mouse versican (GAG*β*) antibody (1:1000) (Millipore-Sigma, catalog no. AB1033) or mouse monoclonal GAPDH antibody (1:5000) (Millipore-Sigma, catalog no. MAB374) were diluted in blocking buffer and incubated overnight at 4°C with gentle rocking. LI-COR IR dye secondary antibodies against mouse and rabbit antibodies (LI-COR Biosciences, Lincoln, NE) were used at 1:10000 dilutions in PBS to detect primary antibodies. A LI-COR Odyssey CLx scanner and the LI-COR Image Studio (ver. 4.0) was used to image western blots.

### Fluorophore-assisted carbohydrate electrophoresis

FACE procedures used were essentially as previously published [4–6]. In brief, tissue was extensively digested with proteinase K and released glycosaminoglycans (GAGs) were isolated by ice-cold ethanol precipitation and centrifugation. GAGs were resuspended in 0.1 M ammonium acetate, pH 7.0 and then digested overnight with *Streptomyces* hyaluronidase (200 mU) at 37° C for HA-FACE. HA digestion products were separated from other intact GAGs using ice-cold ethanol precipitation and centrifugation. The supernatant was then dried under vacuum centrifugation and reacted with 6.25 mM 2-aminoacridone (AMAC) and 625 mM sodium cyanoborohydride (NaBH_3_CN) overnight at 37° C. The pellet was resuspended in 0.1 M ammonium acetate, pH 7.0 and then digested overnight with chondroitinase ABC (25 mU) at 37° C for CS-FACE. CS digestion products were separated from other intact GAGs using ice-cold ethanol precipitation and centrifugation. The supernatant was then dried under vacuum centrifugation and reacted with 6.25 mM AMAC and 625 mM NaBH_3_CN overnight at 37° C. AMAC-labeled HA and CS digestion products were loaded onto 20% (w/v) acrylamide gels and these products were resolved by electrophoresis at 500 V constant current over a 50-60 minute run time. FACE gels were imaged while still housed in the gel plates on a UVP ChemiDoc-It^2^ 515 system. Digital gel images were analyzed for AMAC-labeled HA and CS digestion product band intensities using ImageJ (NIH, Bethesda MD).

### Transmission electron microscopy

E9.5 yolk sacs were harvested from timed pregnancies and fixed in 4% PFA + 2.5% glutaraldehyde in PBS overnight. Samples were dehydrated in an ethanol:PBS gradient and fully dehydrated samples were embedded in pure Eponate12 resin and allowed to polymerize overnight. Ultrathin sections (85 nm) were stained with uranyl acetate and lead citrate and viewed and imaged on a FEI Tecnai G2 Spirit BioTWIN Transmission Electron Microscope (FEI company, Hillsboro, OR) equipped with an Orius 832 CCD camera (Gatan, Inc., Pleasanton, CA).

Some E8.5 embryos were fixed in 4% PFA + 2.5% glutaraldehyde + 0.7% w/v ruthenium hexamine trichloride (Sigma-Aldrich, catalog no. 262005) in PBS overnight to optimize proteoglycan and cell membrane preservation [7] and embedded in Eponate 12 resin as described above. 1 micron thick sections were stained with 1% toluidine blue containing 2% sodium borate for 30 seconds on a hot plate and washed with distilled water. Slides were mounted with Cytoseal XYL mounting media (ThermoFisher Scientific, catalog no. 22-050-262) and imaged as described for routine microscopy.

### VEGF_165_ and Ihh treatment of RPE-1 cells

Wild type hTERT RPE-1 (ATCC, CRL-4000) cells and ADAMTS9 deficient RPE-1 cells [2] were cultured in 8-chamber cell culture slides (Fisher Scientific, catalog no. 354118) with a seeding density of 50,000 cells/chamber in DMEM F-12 culture medium containing 10% FBS for 48 hrs and washed with PBS and cultured in DMEM F-12 culture medium without FBS for an additional 24hrs in a cell culture incubator at 37C with 5% CO_2_. Cells were cultured in DMEM F-12 medium containing 25, 50, 100 ng/ml recombinant human biotinylated VEGF_165_ (R&D Systems, Cat. No. BT293-010) or recombinant human/mouse Ihh N-terminus (C2811) (R&D Systems, Cat. no. 1705-HH-025) for 6hrs. Cells were washed three times with PBS and fixed in 4% PFA. For versican depletion experiments, a previously validated highly potent *VCAN* siRNA (Ambion, Cat. no. S229335) [8–10], or control siRNA (Ambion, cat. no. 4390843) were added to the cell cultures 24hrs after seeding using the Lipofectamine RNAiMAX protocol (Invitrogen, Cat. no. 13778) and cultured for 24hrs prior to serum starvation and growth factor treatment. For treatment with chondroitinase ABC (0.2 U/ml in PBS, Sigma-Aldrich, Cat. no. C3667), cells were incubated overnight prior to the addition of recombinant VEGF_165_ or IHH. For co staining, recombinant VEGF_165_ was detected using Streptavidin FITC (1:400, Invitrogen, SA1001), recombinant IHH using anti mIhh-N antibody (1:400, R&D Systems, Cat. no. AF1705) and versican using the pVC antibody at a 1:400 dilution as previously described [2, 3]. Fixed cell culture chambers were blocked and incubated with primary antibodies in 5% normal goat serum for VEGF_165_ experiments and 5% normal horse serum (Sigma-Aldrich, Cat. no. H0146-10ml) for Ihh experiments. Goat anti rabbit Alexa-568 secondary antibody (Invitrogen, Cat. no. A11011) and donkey anti goat Alexa-488 (Invitrogen, Cat. no. A11055) secondary antibodies were used at 1:600 dilution for detecting versican and Ihh primary antibodies. Super resolution confocal microscopy was conducted as previously described [2].

### Embryoid body formation and VEGF_165_ induced angiogenic-sprouting

mESCs were cultured for two days till 100% confluent, gently washed with PBS and incubated with IMDM without LIF for an additional 48 h. IMDM without LIF was used in subsequent manipulations to form EBs. The medium was removed and 250μl of trypsin was added to each plate and aspirated out after a 30 second incubation leaving a thin film of trypsin. Upon detachment and rounding of cells, 1 mL of IMDM was added to each plate, and a small pipette tip was used to generate small aggregates mechanically from the mESC sheet. The aggregates were transferred to 60 mm (non-adhesive) bacteriological dishes (Corning, catalog no. 351007) containing 5 mL of IMDM, where most curled and formed free-floating spheroids after 24 hrs. These 1-day EBs were transferred to a fresh bacteriological dish with 5 mL of IMDM and cultured for an additional 3 days. 10-20 4-day old EBs from each genotype were transferred to a 4-well chamber (Fisher Scientific, catalog no. 354114) and embedded in bovine type 1 collagen (Corning, catalog no. 354231) containing 30 ng/mL recombinant VEGF_165_ (PreproTech, catalog no. 100-20). 500 μL IMDM containing 30 ng/mL recombinant VEGF_165_, was added atop each collagen gel and replaced very 4 days for a further 12 days. The collagen-embedded EBs were washed with PBS and fixed in 4% PFA for 15 min, permeabilized by three washes in PBS containing 0.3% Tween 20 for 30 minutes prior to immunostaining with anti-CD31 or anti-smooth muscle *α*-actin (SMA). For qRT-PCR, EBs in floating culture were incubated for 10 days (day-10 EBs). 300 μL of TRIzol reagent (ThermoFisher scientific, catalog no. 15596026) was used to harvest total RNA from each 60 mm plate.

### Embryoid body vascular sprout quantifications

For quantifying EB vascular sprouts, collagen-embedded, fixed embryoid bodies in 4-chamber slides were imaged using an inverted bright field microscope. Images were analyzed by Image J-FIJI (NIH, Bethesda, MD), and the freehand drawing tool was used to trace the length of individual vascular sprouts. A total N= 72 wild type, 68 D8 and 68 F9 EBs were analyzed in 3 independent experiments. Lengths of 402 sprouts from wild type, 91 from D8 and 97 from F9 EBs were measured.

## KEY RESOURCES TABLE

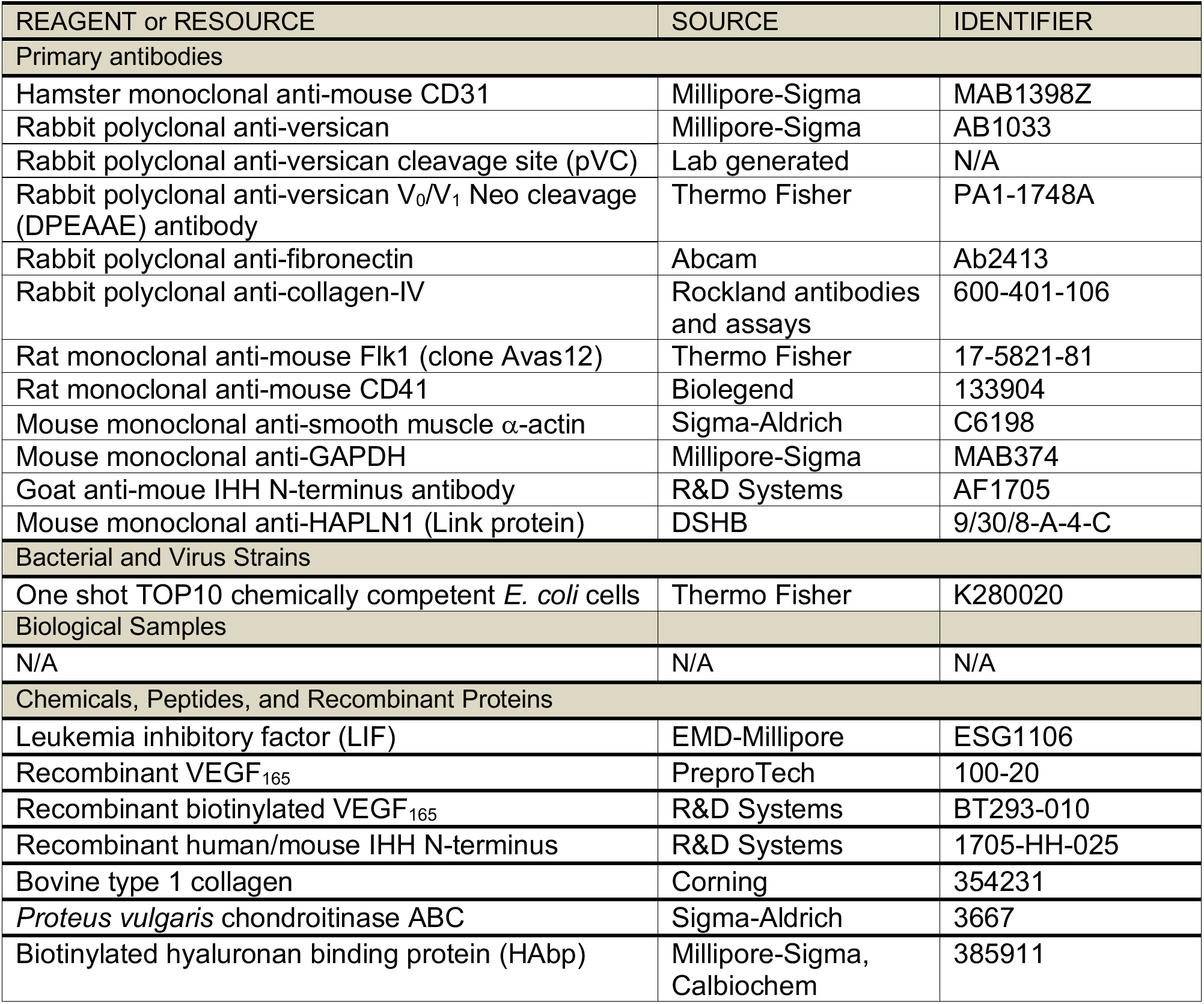
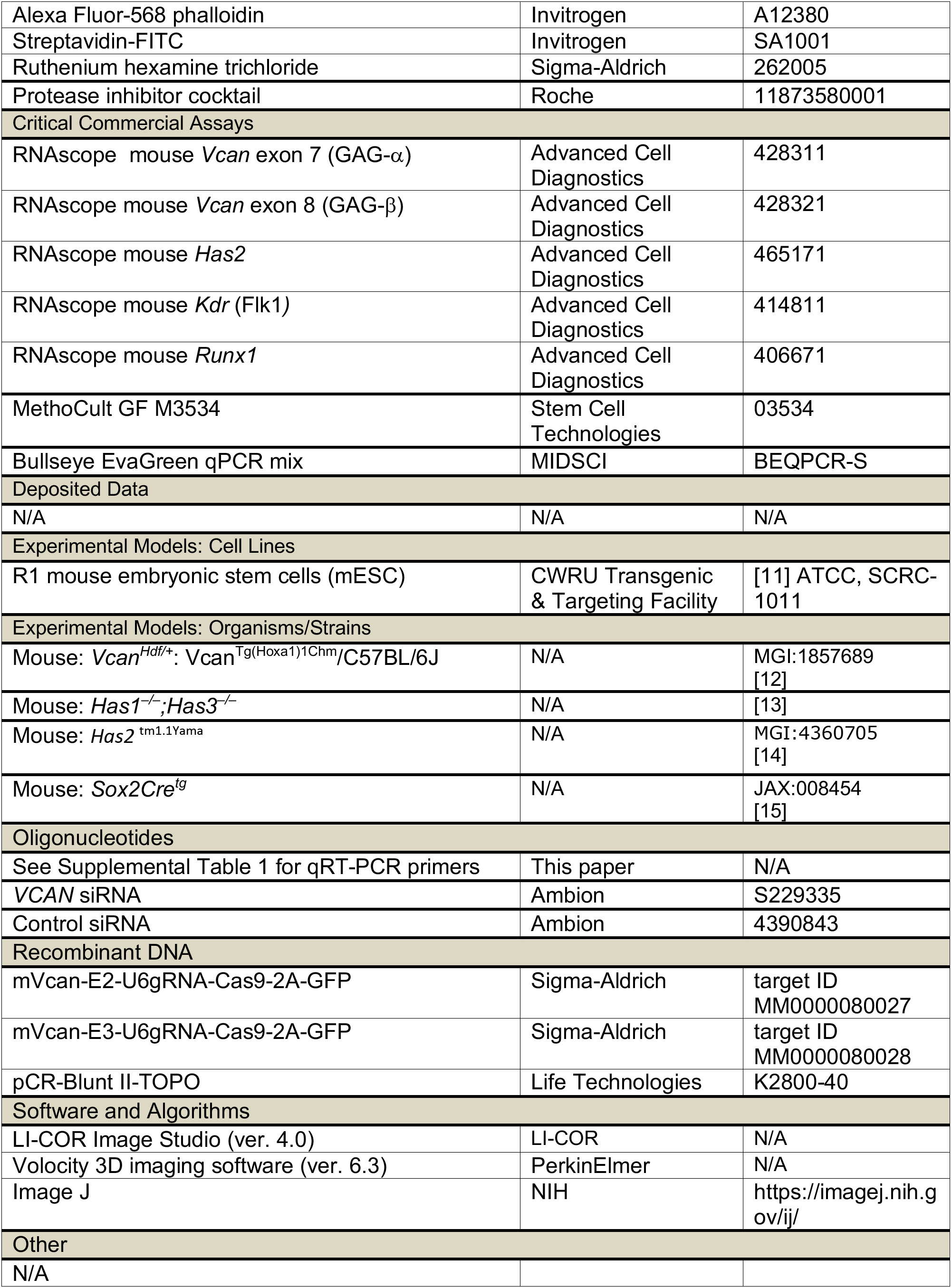

